# Probabilistic Perceptual Decisions During Navigation Are Driven by Small Subpopulation of Neurons in a Single Cortical Column of Primary Somatosensory Cortex

**DOI:** 10.1101/2025.02.17.638764

**Authors:** Alex G. Armstrong, Yurii A. Vlasov

**Affiliations:** University of Illinois Urbana Champaign, Neuroscience Program, Electrical and Computer Engineering, Carle Illinois College of Medicine, Chan Zuckerberg Biohub Chicago

## Abstract

Perceptually driven behavioral choices are thought to develop gradually from sensation to perception in the somatosensory cortex to guide decision-making in higher order cortical areas. Primary somatosensory cortex (wS1) of rodents related to their mystacial whiskers has been a model system to study this information flow. However, the role of wS1 in this process is often debated based on controversial results of loss-of-function behavioral experiments that often require prolonged training and movement restraints. Here, to elucidate the role of wS1 in decision-making, we developed an ethological whisker-guided virtual reality (VR) paradigm that closely mimics natural navigation in underground burrows. Untrained mice navigate left and right turns at high speed by sensing VR walls with just a pair of their C2 whiskers. Inactivating layer 4 of C2 barrel results in loss of ability to produce turns contralateral to the lesion. Using probabilistic model of collision avoidance in the presence of noise and uncertainties we hypothesize that wS1 is involved in a feedback control loop that requires continuous updates and predictions to infer the optimal path for collision avoidance.

**Significance:** Perceptual decisions driven by sensing salient changes in the environment are thought to develop from sensation in primary cortex (S1) to decisions in pre-motor cortical areas. However, the role of S1 in this process is debated based on controversial results of loss-of-function behavioral experiments that often require prolonged training and movement restraints. Here, by utilizing an ethological whisker-guided virtual reality, we show that perceptual decisions causally depend on small subpopulation of neurons in layer 4 of a single cortical barrel. Whisker-guided navigation requires continuous updates and predictions of relative positions of the body and obstacles to infer the optimal path for collision avoidance. These complex computations are likely to rely on nested feedback loops that directly involve wS1 hence making it indispensable.

## Introduction

Perceptual decision-making driven by sensing salient changes in the external environment is thought to gradually develop (de Lafuente and Romo, 2006) from sensation in primary somatosensory cortices (S1) (Hernández et al., 2000) to perception in higher order cortices and further transformed to decisions in pre-motor cortical areas (Roitman and Shadlen, 2002). Rodents’ mystacial vibrissae (whiskers) system with anatomically distinct representation of individual whiskers in the primary whisker-related wS1 (barrel cortex) (Staiger and Petersen, 2020) has been a pivotal model to study this serial information flow (Kleinfeld et al., 2006; von Heimendahl et al., 2007; Feldmeyer et al., 2013; Guo et al., 2014a; Sofroniew et al., 2015; Fassihi et al., 2017; Stüttgen and Schwarz, 2018) that relays extracted sensory features downstream for further processing (Diamond and Toso, 2023). Inactivation of wS1 using lesions (Hutson and Masterton, 1986; Krupa et al., 2001; Chakrabarti and Schwarz, 2018; Ryan et al., 2022; Waiblinger et al., 2022), pharmacological intervention (Krupa et al., 1999), or optogenetic silencing (O’Connor et al., 2010; Sofroniew et al., 2015; Gardères et al., 2024) results in observable changes in behavior and points to the focal role of wS1 in perception and decision making. Inactivation of wS1 degrades performance on whisker passive touch detection (Miyashita and Feldman, 2012), active object location (O’Connor et al., 2010; Guo et al., 2014a), aperture discrimination (Krupa et al., 2001), gap crossing (Hutson and Masterton, 1986; Shih et al., 2013), texture discrimination (Guic-Robles et al., 1992), and whisker-guided navigation (Sofroniew et al., 2015) tasks. This role, however, is frequently debated (Stüttgen and Schwarz, 2018) as many loss-of-function studies(Hutson and Masterton, 1986; Hong et al., 2018; Warren et al., 2021) indicate that wS1 may be dispensable if not for all but at least for some of whisker-dependent behaviors including vibration frequency discrimination (Hutson and Masterton, 1986), passive touch detection (Hong et al., 2018; Waiblinger et al., 2022), and obstacle detection (Warren et al., 2021).

Among many variables that can potentially contribute to these seemingly controversial conclusions, two in particular are of most interest: design of a behavioral task and localization of loss-of-function perturbation. First, studies of decision making typically involve multiple repetitions of stereotypical perceptual tasks that often require prolonged training and movement restraint. Concerns have been raised (Carandini and Churchland, 2013; Krakauer et al., 2017; Pereira et al., 2020; Dennis et al., 2021; Miller et al., 2022; Oesch et al., 2024) that reductionist approaches to behavior task design produce behavioral dynamics far from exercised in natural habitat, thus providing strongly biased dynamics of neural activity that is not necessarily representative of what brain circuits have evolved to produce. Second, localization of loss-of-function perturbation is not always under experimental control and typically affects large neuronal populations that hinders interpretation. Loss-of-function studies typically rely on inactivation of the whole barrel cortex (Hutson and Masterton, 1986; Krupa et al., 2001; Chakrabarti and Schwarz, 2018; Hong et al., 2018; Warren et al., 2021; Ryan et al., 2022; Waiblinger et al., 2022) (cortical volume ∼2mm^3^) with average neuron density (Herculano-Houzel et al., 2013; Keller et al., 2018) of 8.3×10^4^ neurons/mm^3^ that corresponds to 1.7×10^4^ neurons. More localized lesions produced with laser ablation (Ryan et al., 2022) target a single barrel column (cortical volume ∼0.1mm^3^) that contains 8×10^3^ neurons(Lefort et al., 2009). Transient optogenetic (O’Connor et al., 2010; Sofroniew et al., 2015; Gardères et al., 2024) silencing can inactivate smaller volumes, however it often elicits off-target downstream effects (Otchy et al., 2015) affecting multiple brain areas.

To address these challenges, we developed a tactile virtual reality (VR) whisker-guided navigation paradigm (Sofroniew et al., 2014; Sofroniew et al., 2015) that closely mimics high-speed navigation of these nocturnal animals in the dark underground winding burrows of their natural habitat. We show that no prior training or reinforcement is needed for mice to successfully navigate left and right turns in this tactile VR while producing a natural two-beat diagonal trotting gait at high running speeds. This Alternate Navigation Choice (ANC) active behavior is produced even when all but a pair of C2 whiskers are trimmed. To localize loss-of-function perturbation we produced ultra-small electrolytic lesions with volumes below 3×10^-3^mm^3^ using a multielectrode probe array specifically targeting only layer 4 in a single principal whisker barrel column. We show that perceptual decision-making causally depends on a small subpopulation of neurons in layer 4 of a single cortical barrel column. We hypothesize that our untrained and unrewarded behavioral paradigm requires continuous probabilistic predictions on relative positions of the animal body and VR walls even in the presence of large sensory noise and obscured sampling. Therefore, computational resources needed for continuous wall tracking may significantly exceed those required for tasks where appearance of an obstacle along the navigation path is reported just once and collision avoidance planning may not involve barrel cortex (Hong et al., 2018; Warren et al., 2021).

## Materials and Methods

### Animal preparation

#### Datasets

Thirty-three mice aged between P63-P115 of both sexes were used for these experiments, with 7 used for behavioral experiments with all whiskers intact and 26 used for experiments where their whiskers were trimmed so just a pair of C2 whiskers remained (Table S1), with 15 of these mice were used for lesion tests, with 7 undergoing electrophysiological recordings. To ease identification of principal barrel and enable optogenetic tagging, the transgenic *Scnn1a-TG3-Cre* (Jackson: 009613) line was crossed with the *Ai32* line (Jackson: 012569), producing strong co-expression of enhanced yellow fluorescent protein (EYFP) and Channelrhodopsin-2 (ChR2H134R-EYFP) in ∼85% of layer 4 excitatory neurons in the primary somatosensory cortex(Madisen et al., 2012). All procedures were in accordance with protocols approved by the University of Illinois Urbana-Champaign Institutional Animal Care and Use Committee (*protocols #20138, #23128*). Animals were kept on a reverse 12-hour light/dark cycle in individual cages equipped with activity wheels (K3250 Fast-Trac, Bio-Serv) to encourage active behavior in-between experimental sessions. All experimental sessions were performed during the dark phase.

### Experimental Design and Statistical Analysis

#### Headbar implantation

Surgical procedures were carried out aseptically while animals are anesthetized with isoflurane (1-3% in oxygen) before being injected subcutaneously with 5mg/kg carprofen. To prepare animals for behavioral and electrophysiological studies a custom-built titanium headbar (10mm x 2 mm)(Guo et al., 2014b) was attached to the animal’s skull to hold the animal’s head using Vetbond (3M, USA). Orientation of the headbar is carefully adjusted with respect to Bregma and Lambda points on the animal skull to provide a repeatable coordinate system for multielectrode probe insertion. The central open area of the headbar (3mm x 2mm) exposes the skull directly above the primary somatosensory cortex for in-vivo visualization, targeted craniotomy, and precise probe insertion. This area is covered in optical grade clear cement for in-vivo fluorescence imaging. For experiments with electrophysiological recordings, a craniotomy (approximately 0.2 mm in diameter) is made through the back of the headbar, and a ground wire (2 mm long Pt/Ir ground wire, A-M Systems 776000) is implanted in the cerebellum and cemented on the headbar. Beginning the day following the surgery, subcutaneous Carprofen (5mg/kg) is administered for two days to reduce inflammation.

#### Whisker trimming

Mice in the single whisker cohort have all whiskers apart from C2 on either side trimmed. Mice are anesthetized with isoflurane (1-3% in oxygen) before, with trimming repeated at least every 2 days to prohibit other whiskers from growing back. Whiskers are first trimmed on the first day of animal’s first behavioral session. This is to reduce the time spent with just a single whisker to ≤6 days. In adult mice, morphological changes associated with whisker trimming take at least two weeks to form(Fox, 2002).

### Behavior Rig

#### Tactile VR experimental rig

The treadmill of **Fig.1A** is built around a hollow Styrofoam sphere (16-inches diameter) with walls custom-thinned to provide a total weight of less than 60g. The sphere is suspended on 7 ping-pong balls supported by 7 air-nozzles distributed around its the bottom half based on the design developed at the HHMI Janelia Research campus. Controlled stimulus to animal whiskers is provided by a pair of motorized walls positioned at both sides of the snout. The animal movement in lateral (V_lat_) and forward (V_for_) directions is tracked at 500Hz by recording the sphere movement with two tracking cameras. Turns were defined by a turn angle *θ* that was positive for left turns and negative for right.

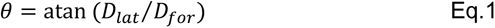

where *D_for_* and *D_lat_* are lateral and forward displacements in a single cycle. This signal is used to provide positive feedback for the movement of the walls (V_wall_):

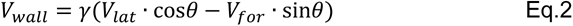

where *γ* is a calibrated feedback coefficient. When taken as negative it generates a positive closed-loop (CL) virtual reality (VR) that moves walls closer to the snout if the animal is moving closer to them and, therefore, encourages the animal to stay in the middle of the VR corridor.

**Figure 1.**
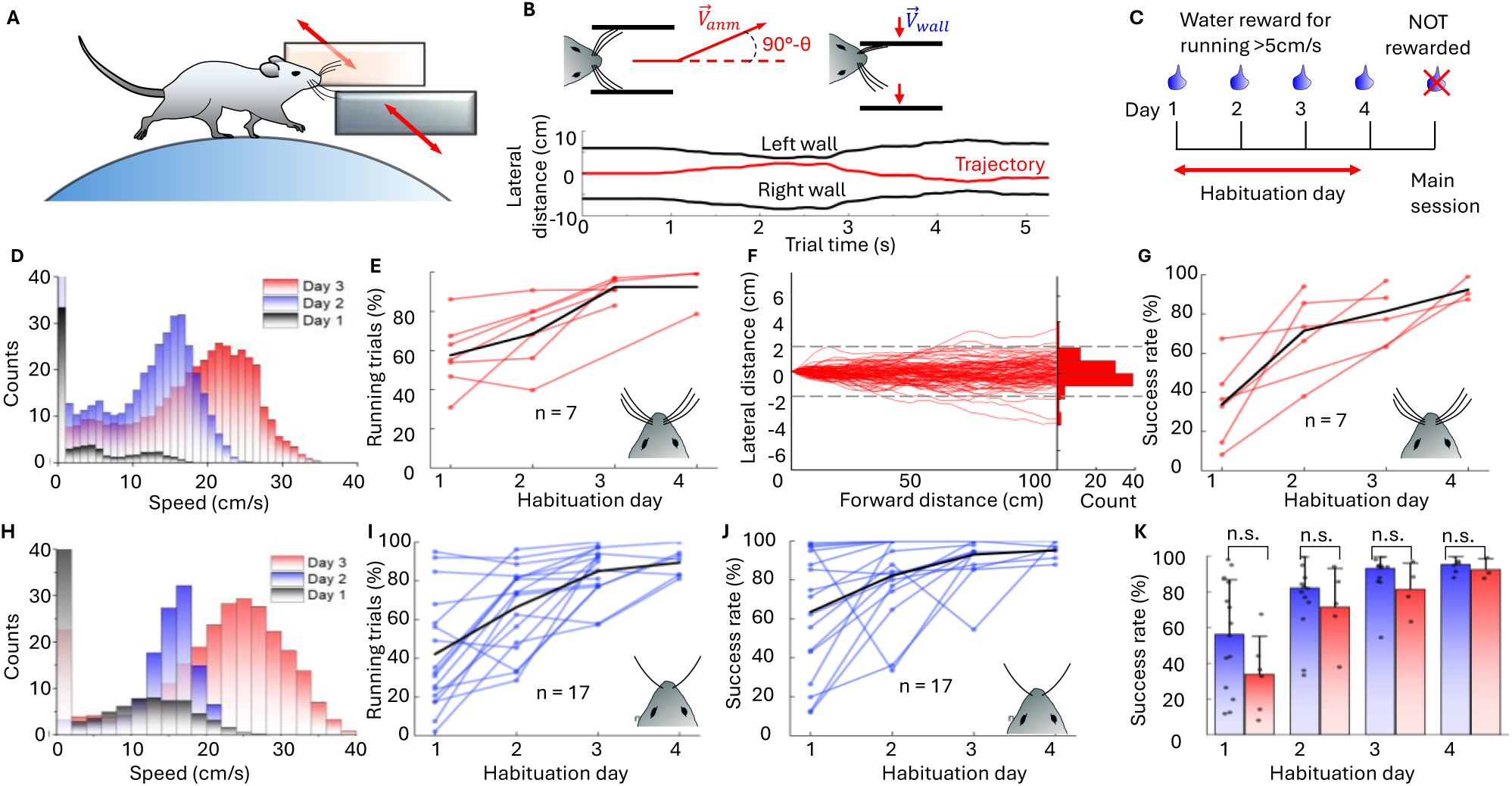
High-speed tactile VR navigation tasks are untrained, unrewarded and can be performed with just a pair of C2 whiskers. **A)** Schematic of the tactile VR rig with mouse head-fixed on air-suspended sphere. Two motorized walls on either side of the snout are the only sensory cues for mice to track with whiskers and navigate. **B)** Top: Closed-loop control of wall movement coupled to the mouse movement trajectory (Eq.1). Bottom: example 5 second segment of a straight run CL trial. When an animal is moving closer to the left wall, the left wall is moving closer to the snout encouraging animal to move closer to the center of VR corridor. **C)** Experimental protocol includes 3-4 days of habituation to CL navigation in VR as shown in B) when animal is rewarded for running at elevated speeds. This is followed by a single main session lasting for up to 2 hours that is unrewarded to quantify performance and introduce ANC tasks. **D)** Histograms of speed distribution across 3 habituation sessions for a single animal with all whiskers intact. **E)** Percentage of time in each habituation session where mice with all whiskers (n=7) are running faster than 5cm/s. **F)** Left: Trajectories of a mouse during a single main session with CL straight run. Each 100 overlaid individual trials correspond to 100cm forward distance. Success threshold shown by dashed lines with ±2cm clearance. Right: Histogram of lateral distance for 100 trials showing over 90% success rate. **G)** Success rate during 4 habituation sessions for the same 7 mice. Black line shows mean across animals. **H)** Histograms of speed distribution across 3 habituation sessions for a single animal with all but C2 whiskers trimmed. **I)** Percentage of time in each habituation session where mice with just a pair of C2 whiskers (n=17) are running faster than 5cm/s. **J)** Success rate during 4 habituation sessions for the same 17 mice. Black line shows mean across animals. **K)** Statistical comparison of success rate over training sessions between mice with all whiskers intact (blue) and mice with just a pair of C2 whiskers (red). two sample *t*-test, day 1 p = 0.083, day 2 p = 0.363, day 3 p = 0.212, day 4 p = 0.519.

The real time (RT) control and acquisition system is custom designed to power a FPGA CompactRIO (cRIO-9045, NI) with 32 channel analog voltage input module (NI 9205) to read sphere velocities, 4 channel analog output module (NI-9263) to operate mechanical walls, and 32 channel TTL input/output (NI-9403) to provide synchronization to video cameras and other equipment. The module is operated via LabVIEW RT Module (NI) by the custom designed LabVIEW acquisition program that has Master/Slave architecture. The program operates several parallel processes that need to run simultaneously and continuously but at different rates. The master process runs in a loop at 30MHz and provides RT analog acquisition and control data to the rig. The master loop controls several slave loops and communicates with them using messaging architectures. One of the slave loops buffers recorded data and logs the data to disk at its own much slower pace. As a result, the acquisition and control experimental protocol executed in the master loop can run continuously without interruptions with virtually no inter-trial interval. Another slave loop is designed to provide a graphical user interface (GUI) that allows the user to observe data on the screen such as overlaid trial trajectories of the animal and to change parameters for each trial session if needed. Control signals generated in the master loop are directed to synchronize 3 high-speed cameras (**Fig.1A, Video S1**) to acquire videos of whiskers and paws, as well as to synchronize behavioral recordings with electrophysiological recordings, optogenetic stimulation, and provide water rewards to the motorized licking port if needed.

The VR rig is enclosed in a light-tight box with fine copper mesh on the inside to provide shielding from ambient light and EM interference. Constant hissing sound from air-nozzles provides stationary background white noise that masks all other ambient noises in the room. The surface of the Styrofoam sphere is coated with permanent water-resistive protective coating using low-odor latex aerosol (Krylon 7120) to enable the sphere to be cleaned between each animal thus minimizing olfactory cues. Therefore, the only sensory cues left for animals to use to navigate in the VR are whiskers palpating the motorized walls. The full LabVIEW program is available on our GitHub page (https://github.com/NeuroTechnologies).

#### Habituation

Following full and complete recovery from a previous surgery, the animal is placed on a water restriction schedule in preparation for behavioral conditioning. Food is continuously available. Water is adjusted (approximately 1 ml per day) so that mice maintain more than 80% of free-drinking weight. When fully adjusted to the water restriction protocol (weight stabilized) a mouse is habituated to virtual reality treadmill (**Fig.1A)** with head fixation starting with ∼15min during the first day. Using a large diameter sphere (16 inches) in our VR rig as well as tunable head fixation closely mimics the posture observed during natural locomotion. After the first day, sessions are extended over 3-4 days to last up to 1 hour where the walls are introduced in closed loop straight running trials of increasing distances. During this time a mouse is encouraged to run by administering water rewards (0.02ml) given only if the running speed exceeds 5cm/s for a total of 1 ml. Typically on day 3 a mouse is fully acclimatized to the apparatus, actively runs 80% of the time at elevated speed above 5cm/s and consumes all the daily water during the experiment. When running, they navigate the straight VR corridor while minimizing the lateral displacement to less than ±2cm. We regard the animal as habituated when running ≥75% of the session, with ≥80% within success criteria (when running) at the maximum trial distance of 100cm, with a ≥50 trials of this type.

#### Trial structure

For main sessions, trials consisted of two epochs, 100cm of baseline closed loop VR straight running where walls are positioned mean 10mm from either whisker pad, followed by open loop turn to the left or right lasting for 2s. Walls are decoupled and one wall is moved closer to the whisker pad of the mouse (2mm closest distance), while the other is moved out of reach. The oddball stimulus requires the mouse to turn in the VR environment away from the wall. Success criterion was defined as whether the animal has a mean run angle change of at least 10°. The main session is completely unrewarded and has no other sensory cues.

#### Whisker tracking and reconstruction of whisker angle and curvature

High speed overhead video of the whisker (**Video S1**) was captured simultaneously with behavioral recordings using the overhead 656 × 600 pixels camera at 1000–1000 fps (EoSens1, Mikrotron) equipped with a 0.36× telecentric lens that produced 25 mm × 23 mm field-of-view (Edmund Optics, no. 58-257). Video streams were digitized with a frame grabber (BitFlow Axion) controlled by StreamPix7 multicamera software (NorPix). To align videoframes to the master clock with less than 1 ms jitter, a video system was triggered from the main RT LabVIEW acquisition program. The field of view (FOV) is adjusted (MC ControlTool, Mikrotron) to capture the full length of the C2 whisker as well as its bent shape when interacting with a wall. The whisker was illuminated by an overhead LED light source (ThorLabs M810L3) powered by DC power supply (Mean Well RS-15-24) and focused with 40 mm focal length aspheric condenser (Thorlabs, ACL5040U).

Whisker movement during interaction with the wall was tracked by extracting individual lossless frames for subsequent manual analysis or using automated tracking with SLEAP (Pereira et al., 2022) model. A skeleton of five points along the C2 whisker was manually identified for 50-100 frames for each video and used as the training dataset for the model. Once trained, the model was used to make inferences on all frames in the video using the Hungarian matching method. The angle of the whisker is defined as 0° when it is perpendicular to the mouse head AP axis. The first two tracking points along the whisker length in each frame were used to calculate the whisker angle. To extract whisker curvature from the tracking results, a circle was fit to the three points closest to the base of the whisker and curvature defined as inverse of the radius.

#### Simultaneous tracking of fore and hind limbs and reconstruction of gait

Videos of hind paws and fore paws were captured using a pair of 659 × 494 pixels cameras at 50fps (Basler acA640) equipped with 16 mm lenses that produced 34 mm × 23 mm FOV. Cameras were positioned above the hind limbs and to the side of the front limbs (**Fig.1A**) and were triggered from the main RT LabVIEW acquisition program. The FOVs were illuminated by additional LED light sources. Frame-by-frame manual inspection of the paw’s videos enabled reconstruction of the animal gait (**Video S1**) by recording the precise timings of when each paw is in contact with the sphere. This reconstruction is confirmed by observation of characteristic periodic oscillations (**Fig.3B**, middle) in the recorded traces of turn angle *θ* derivative (or lateral acceleration). To classify navigation periods as either trotting or walking, wavelet analysis was performed on the lateral acceleration trace, showing lower frequencies (<4Hz) were present in only walking trials and absent during trotting (**Fig.S1C**). Step frequency and mean speed were calculated for each trial and a Fisher’s linear discriminant analysis (LDA) was performed to classify segments as either walking or trotting (**Fig.S1B**). This classifier was applied to all animals with a mean 6% error rate after manual assessment (**Fig.S1D**). Due to mice switching gaits between trials, only those with a probability of walking or trotting >0.8 were used for comparisons.

### Lesion studies

#### Stereotaxic-less probe positioning on a principal whisker barrel

We aim for precise placement of microelectrode probe strategically into the principal whisker barrel and to align it perpendicular to cortical layers whilst the animal is placed and head fixed in the VR rig. Therefore, we developed a protocol for highly precise probe insertion without using stereotaxic equipment.

The first step is to acquire a brain map of the somatosensory cortex generalized across many animals (**Fig.5C**). For that we used fluorescence top-down imaging (Olympus, MVX10) of the whole fixed brain harvested immediately after the final terminal experiment (**Fig.5B**). To match vasculature patterns with those observed in-vivo in the same animal (**Fig.5A**), mice were deeply anesthetized with 5% isoflurane then perfused with 0.1 M sodium phosphate buffer for just a short period of time to ensure that some blood remains in top arterioles on the brain surface. Resulting brain preparations not only show characteristic contours of primary visual cortex (V1), the barrel cortex (wS1), as well as characteristic shapes of lower jaw (LJ) and hind paw (HP) somatosensory areas, but also in some cases show identifiable locations of main barrels across main B, C, and D rows. Combining similar images from several animals, a generalized map is created with positions of individual barrels relative to the contours of V1 and S1 areas (**Fig.5C**).

To map in-vivo the location of the target C2 barrel without the use of stereotaxic apparatus, the mouse is anesthetized with isoflurane (2-4% in oxygen) and the brain is imaged in-vivo (**Fig.5A**) through the central open area of the headbar that exposes the skull directly above the primary somatosensory cortex (nominal coordinates for C2 barrel 2.0mm AP, 3.4mm ML). The YFP expression in L4 excitatory cells in the primary somatosensory and visual cortices is strong enough to identify contours of V1, wS1, LJ and HP areas as well as characteristic vasculature patterns through the skull, and to align them to previously acquired cortical map (**Fig.5B**). This procedure allows us to mark the location of the C2 barrel with respect to the headbar with localization precision as high as ±0.1mm as confirmed by subsequent histological mapping. For targeting of neighboring barrel, final position was adjusted by ∼200µm.

#### Targeted craniotomy

On the morning of the final terminal experimental lesion session, mice are briefly anesthetized with isoflurane (2-4% in oxygen) and are placed in a head-fixation surgery apparatus closely mimicking head-fixation setup in the VR rig. Fluorescence imaging is performed with overhead stereo microscope (MZ12.5, Leica) equipped with fluorescence adaptor (Quad Fluorescence System, Kramer Scientific). The brain map with location of targeted C2 principal whisker barrel generated during in-vivo fluorescent imaging is compared with microscope imaging to check the marked location of targeted craniotomy on the skull surface. Using a microdrill with 0.002” burrs (Fine Science Tools, Canada), the optical cement and the skull are carefully thinned while the surface is cooled down using sterile artificial cerebrospinal fluid (ACSF). A small craniotomy of < 300µm diameter is opened while the dura remained intact. The craniotomy site is then imaged both in a white light and with fluorescence filters to visualize a barrel field and a vasculature pattern and to produce a final map for probe implantation. The craniotomy site is covered with a silicone elastomer plug (World Precision Instruments, USA). The whole procedure typically lasts less than 30 mins and mice are returned to home-cage for recovery for at least 2 hours. Typically, mice are fully recovered within 20 mins and frequently run on activity wheels.

#### Targeted lesions

To localize inactivation to the main thalamocortical input layer (L4) of a barrel column, 64 channel linear shank silicone probes (Cambridge NeuroTech, UK) were used. Connected to an Intan headstage (Intan, USA) via a principal component board, an external wire was soldered to the pin on the PCB that corresponds to the deepest channel of the probe. The probe is inserted into the brain at a 38° angle to be parallel to the barrel column for a total ∼400-500µm depth at a rate of 80µm/min. Probe is left to settle for 5 minutes before optical tagging is done using a 473nm laser (ThorLabs, S1FC473MM) positioned above the barrel of interest. Resulting activity in local field potential is then visualized as a current source density plot (Fig.5A). A stereotypical sink originating from layer 4 spiny stellate cells expressing channel rhodopsin 2 can then be mapped to which electrode on the shank of the probe corresponds to the center of L4. The probe is then advanced or retracted if needed. Once ready to perform the lesion, a constant current isolated stimulator is connected (Digitimer Ltd, UK) while the head stage was disconnected. Then 1mC charge was sent through the wire to the lesion channel, creating the lesion.

#### Lesion quantification

After lesion study was completed, the brain was removed and drop-fixed in 4% paraformaldehyde. Coronal sections 50µm thick were taken using vibratome and stained with DAPI to identify cell bodies (Thermo Fisher, USA). Fifty Z-stack images were taken per slice using Leica SP8 confocal microscope (Leica, Germany). Barrel column layer 4 and lesion volume were accurately measured from reconstruction of these images using 3D Slicer (slicer.org) as well as the same barrel layer 4 in the un-lesioned hemisphere for reference. DAPI staining allowed cell counting within the barrel and lesion itself, confirming that about 10% of the L4 barrel volume containing on average a few hundreds of neurons was inactivated.

#### Behavior-lesion studies

Eight mice were used for lesion studies. Mice were trained the same as all animals with only a single C2 whisker is present on either side. For the main session, mice are running freely on the ball while the probe is positioned and inserted into either the C2 barrel column (n = 4) or a neighboring barrel column (n = 4) and the electrode corresponding to the lesion channel is positioned in the center of layer 4. Once the probe is in its final position the first main behavioral session started, lasting for 10-20 minutes. The session is stopped, the lesion is carried out and the second behavioral session, consisting of the same trial types is started. Total duration between the first and second behavioral session is ∼75 seconds. Due to the lesion always being performed in the left hemisphere, the open-loop turning trials to the left or right correspond to stimulation that is either contralateral or ipsilateral to the lesion site respectively.

#### Electrophysiological recordings

Seven mice underwent electrophysiology recordings before and after lesion experiments following established protocol (Armstrong and Vlasov, 2025). For these animals, the probe was initially inserted 1mm into the cortex and a 20-minute recording was carried out during the pre-lesion behavioral session. Optical tagging was subsequently performed, and the probe was retracted so that the wired lesion channel was in the center of layer 4. The lesion was then created at the same power as detailed in Behavior-lesion studies section, and the probe was readvanced to 1mm depth again to cover all cortical layers. A second 20-minute electrophysiology recording was then carried out during the post-lesion behavioral session. Spikes were sorted using Kilosort 4 (Pachitariu et al., 2024) with improved drift correction to counteract the effects of reinsertion. Metrics were obtained for each unit using Ecephys analysis package (Siegle et al., 2021) and only units that automatically pass standardized quality metric thresholds were included in analysis (presence ratio > 0.9, ISI violations < 0.5 and amplitude cutoff <0.1). From the seven animals, 197 units pre-lesion and 179 units post-lesion passed these thresholds. For spike rate analysis, population activity for each animal was normalized on the number of neurons for each recording, both pre and post lesion.

## Code/Software

### Probabilistic time to collision

#### Deterministic 2D Time to collision

To calculate TTC in 2 dimensions, 3 different scenarios when the collision is with RH paw, RF paw, or nose are considered (**Fig.7A**). For each, a rectangle shape is used whose width is defined by the distance between hind legs (1.8cm), between forelegs (1.5cm) and the width of the snout tip (0.5cm). The rectangle length is equal to the body length (8.4cm from the nose to the tail base for **Fig.7A** schematics). The centerline of the rectangle is aligned with the animal velocity vector 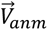. The wall is also represented as a rectangle with a width of 0.2cm and length of 1cm, that can move with 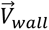 only along lateral direction. Given the state 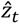 of the animal body (current position, velocity, and acceleration) and corresponding state of the wall, the 2D TTC is calculated as shown in **Fig.7B**.

#### Probabilistic 2D Time to collision

Standard Kalman filter with 4 stages (**Fig.7D**) was implemented in Python. To emulate this internal decision feedback loop that requires measurements (wall via whisking and body state via proprioception) followed by predictions of future relative positions via internal brain models, we use a clock cycle of 8ms to roughly mimic the duration of cortical feedback loops. As a ground truth (GT) we used a 0.3s section from a ANC experiment with recorded positions of the wall and a sphere during OL left turn (**Fig.7E**, dashed black curves). Recorded whisker curvatures and wall contact times are presented in **Fig.S5**, red. Animal is trotting during this section, so we tracked only RH legs for clarity. Corresponding contact times of RH paw with the sphere and angular velocity traces are in **Fig.S5**, blue. For this trial the first whisker wall touch event is at 7ms and the decision to turn is implemented by RH propulsion at the end of a stance at 0.275s that produce characteristic positive peak in angular velocity trace.

We assume that measurements of the wall and body parts positions are available only during their contact times sampled at the clock frequency. Initial state vectors of mouse and wall trajectories were set to zero. We introduced wall position measurement noise by randomly sampling from a gaussian distribution with 1cm SD around the mean taken from recorded GT position. Similarly, body position (centerline) is also taken from recorded GT with additional measurement noise with SD of 0.2cm. We added a process noise due to noise in brain computations and uncertainties in the internal models within standard range for the covariance matrix (0.001-0.1). Probabilistic 2D TTC was calculated using the positional predictions outputs by the Kalman filter for both the body and the wall and subsequent calculations of vector states of both objects. Distribution of TTC values was calculated during each cycle of the Kalman filter, with 20 TTC values produced.

## Results

### Navigation at high-speed locomotion in a tactile VR is successful even when untrained and unrewarded

Virtual reality (VR) has been developed mostly to study visual sensory processing with the movement of a projected imagery scene controlled by animal movements to study dynamics of hippocampal place cells (Dombeck et al., 2010) and visual perceptual decision making (Krumin et al., 2018; Pinto et al., 2018). Tactile VR based on whisker stimulation using a rotating textured cylinder with rotation speed proportional to the animal velocity has been explored to study texture discrimination(Isett and Feldman, 2020), sensorimotor integration (Ayaz et al., 2019), and detection of unexpected sensory signals (Chinta and Pluta, 2023). While such VR generates an illusion of moving textures, animals reported their choices through licking, which typically requires prolonged training (Isett and Feldman, 2020).

To generate a behavioral paradigm in which untrained and unrewarded animal can report its choices naturally, we developed a tactile VR paradigm (Sofroniew et al., 2014; Sofroniew et al., 2015) that engages the animal to choose between left and right turns in response to the movement of motorized walls palpated by animals’ whiskers. Mice are head-fixed on a hollow polystyrene sphere (16-inch diameter, 60g) that is suspended by compressed air, thus acting as a 360° treadmill, with two walls positioned either side of the snout so they are simultaneously reachable with left and right whiskers (**Fig.1A**). Animal’s locomotion 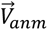 in lateral (*V_lat_*) and forward (*V_for_*) directions is tracked at 500Hz by recording the sphere movement with two tracking cameras that provide feedback to the lateral movement of the walls as described by Eq.2 above. This provides a closed-loop (CL) VR that moves the walls closer to the snout if the animal is moving closer to them and, therefore, encourages the animal to stay in the middle, thus creating a virtual corridor for them to navigate through (**Fig.1B**). The VR rig is enclosed in a light-tight box, broadband sound from air-suspension masks ambient noise and the sphere is cleaned out after each session to remove odors, so that the only sensory cues left for the animals to use for navigation in the VR are the whiskers palpating the moving walls. Thus, our tactile VR closely mimics the navigation in dark burrows of mice in their natural habitat.

Typically, animals require only 3-4 days of habituation sessions with the CL straight trials (**Fig.1C**) to acclimatize to head-fixed locomotion (Methods) with a water reward given only to encourage running at absolute speeds above 5cm/s. During the habituation period mice start to run for longer times with most of them (n=7 in **Fig.1E**) above 75% of the session on day 3 (**Table S1**). While running, they cover total distances approaching 1.5km (**Fig.S1A**); active behavior that is drastically different from typical behavior tasks when animal movement is severely restricted (Hutson and Masterton, 1986; Krupa et al., 1999; Krupa et al., 2001; von Heimendahl et al., 2007; O’Connor et al., 2010; Miyashita and Feldman, 2012; Feldmeyer et al., 2013; Guo et al., 2014a; Burgess et al., 2017; Fassihi et al., 2017; Chakrabarti and Schwarz, 2018; Hong et al., 2018; Odoemene et al., 2018; Ryan et al., 2022; Waiblinger et al., 2022; Diamond and Toso, 2023; Gardères et al., 2024). During habituation mice run at progressively higher speeds as shown for an example animal in **Fig.1D** with its speed increasing from 14.3 ± 4.1cm/s (mean ± s.d.) on day 2 (blue) to 24.3 ± 6.1cm/s (mean ± s.d.) on day 3.

Once habituation is completed, the main completely unrewarded session is performed for 2-hours, with navigation-based decisions introduced (**Fig.1C**). Using only the information from their whiskers interacting with the walls, mice position themselves in the middle of the virtual corridor, which can be quantified using a lateral displacement threshold of ±2cm for every 100cm forward distance (**Fig.1F**). In stark contrast to learned whisker discrimination or decision-making behaviors where reports are given by licking or nose poking that often require weeks-long training, mice in our active VR take just 3-4 short habituation sessions to reach 80% success rate (**Fig.1G**). Notably, once mice are habituated to run head-fixed (**Fig.1E**) at progressively high speeds (**Fig.1D**) they are constantly whisking as running and whisking are strongly coupled (Sofroniew et al., 2014). Therefore, an increasingly high success rate (**Fig.1G**) is unlikely to be the result of learning the whisker-guided navigation, but rather a manifestation of an ability to run head-fixed on a treadmill. Therefore whisker-guided navigation in our VR is likely not learned, but rather a “spontaneous”(Oesch et al., 2024) or “naturalistic” (Sofroniew et al., 2014) behavior.

### High-speed tactile VR navigation tasks can be performed with just a pair of C2 whiskers

To restrict the sensory information flow into the primary somatosensory cortex and localize the cortical circuits for loss-of-function intervention, all but just a pair of the C2 whiskers (second longest mystacial vibrissae oriented almost parallel to the ground) are trimmed. For mice in this cohort (n=17, **Table S1**), whiskers are trimmed on day 1 of the protocol, when they are first placed on the VR rig and retrimmed every day after. Over the same 3-4 habituation sessions, these mice run for longer durations and at higher speeds until they run ≥75% of the session by day 4 (**Fig.1H,I**). Similarly to mice with all whiskers intact, mice with just a pair of C2 whiskers perform the straight run CL task at ≥80% success rate at the end of habituation despite being unrewarded (**Fig.1J**). Note that anatomical barrel field maps are fixed(Feldmeyer et al., 2013) in mature adult animals used in our work (P85 ± 15, mean ± s.d., **Table S1**) with plasticity in response to whisker sensory deprivation building up typically at timescales longer (Staiger and Petersen, 2020) than our experiments (**Fig.1C**). Hence, the high success rate achieved by animals with just C2 whiskers is unlikely a learned, but rather a “spontaneous” (Oesch et al., 2024) or “naturalistic” (Sofroniew et al., 2014) behavior as for animals with all whiskers present. Indeed, statistical analysis shows no significant difference (two-sample t-test, p>0.05) of success rate between the two cohorts of mice not only at the end of the habituation period but also throughout habituation (**Fig.1K**).

### Whisker curvature provides continuous sensory input during VR navigation

Traditionally whisking in rodents is studied in paradigms involving interaction with a pole or other stationary objects that cause whisker deflections (O’Connor et al., 2010). To understand how whiskers interact with the moving walls during whisker-guided navigation and thus what is the sensory information available for the mouse to make navigation decisions, a high-speed overhead camera filmed a C2 whisker at 1000 fps (**Fig.2A, B, C**) while mice were navigating with C2 whiskers during the main behavioral session after completing the habitation. Whisker shape was extracted (Methods) to compute whisker angles, phases, curvatures, and time of wall contacts (**Fig.2D, E, F**). There are three main epochs to consider: epoch t0 (**Fig.2A, D**) of free whisking when the wall is beyond the whisker reach, epoch t1 (**Fig.2B, E**) when whisker is tracking the wall in a series of short palpations, and epoch t2 (**Fig.2C, F**) when whisker is strongly bent when the wall is at its closest position to the snout. Free whisking during epoch t0 (**Fig.2A, D**) is executed at 17.0Hz ± 1.6 (mean± s.d.) with an amplitude of 41° ± 10 (mean ± s.d.) around a set point of 53° ± 3 (mean ± s.d.) that corresponds to a “foveal” whisking, generated at high speed locomotion(Kleinfeld et al., 2006; Sofroniew et al., 2014). Whisker curvature fluctuates around the baseline 3·10^-3^/mm corresponding to natural whisker curvature (**Fig.2D**). During epoch t1 (**Fig.2B, E**) when the wall is moving closer within the whisking range, a whisker periodically palpates the surface with set point (40.4° ± 3.9, mean ± s.d.), amplitude (44.9° ± 13.5, mean ± s.d.), and frequency (17.8 Hz ±1.7, mean ± s.d.) remaining statistically similar (**Fig.2G, H, J).** The curvature changes rapidly during the moments of the whisker touch (**Fig.2E**) occurring mostly during retraction cycles with 44.9% ± 13.4 (mean ± s.d.) of time being in contact with the wall surface (**Fig.2I**). When the wall is at close proximity during epoch t2 (**Fig.2C, F**), the whisker is almost always (84.8% ± 9.6, mean ± s.d.) in contact with the surface (**Fig.2I**). Correspondingly, whisking amplitude is significantly reduced to 17.9° ± 5.4 (mean ± s.d.) and the whisking angle is clamped at 62° ± 2 (mean ± s.d.). However, the curvature is still strongly modulated (**Fig.2F**) at the whisking frequency statistically indistinguishable from epoch t1 and t2 (**Fig.2J**). Therefore, continuous sensory input during navigation is available even when whisking is severely restricted by nearby obstacles.

**Figure 2.**
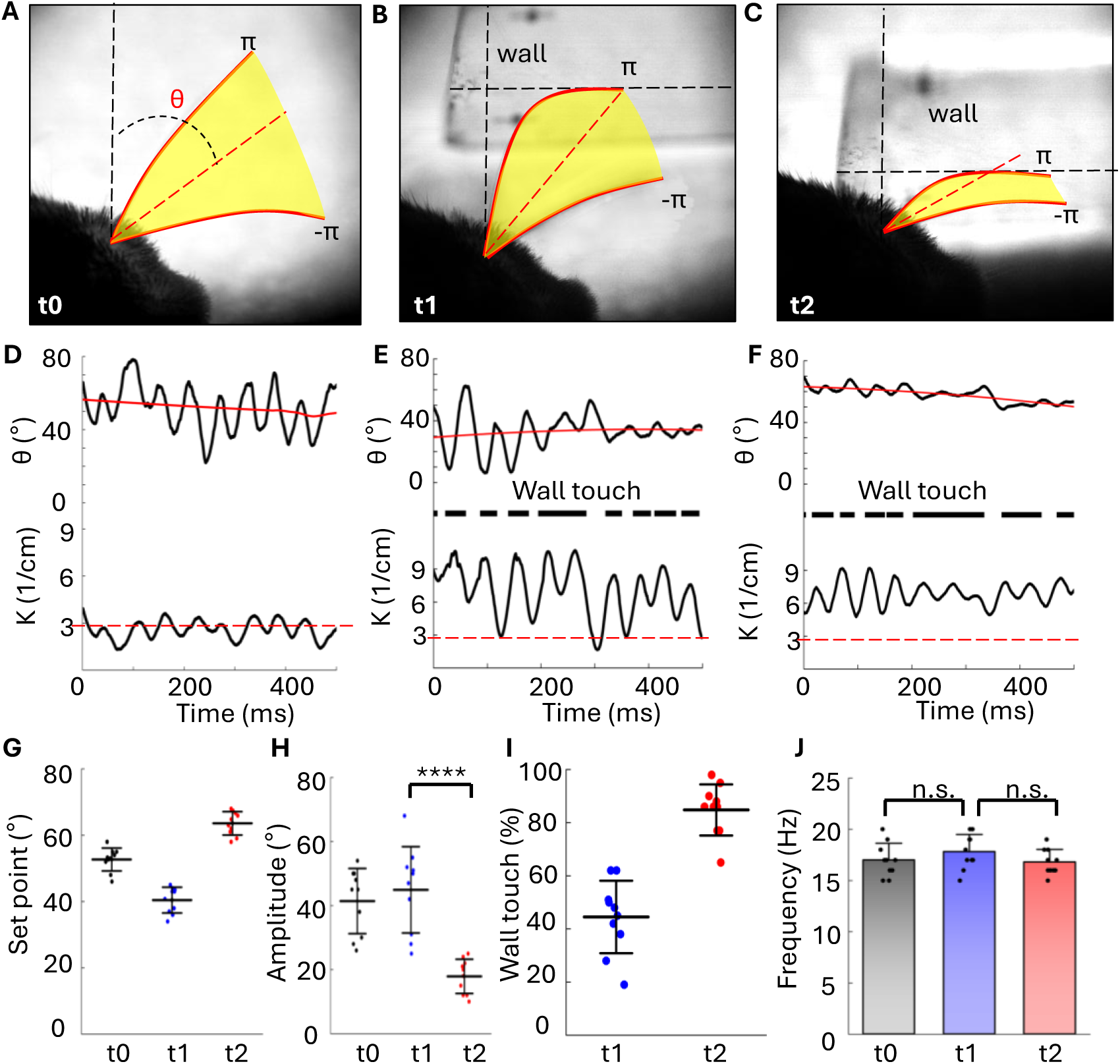
Whisker curvature provides continuous sensory input during VR navigation. **A)** Overlaid video frames of a single retraction cycle of free whisking during epoch t0. **B)** Same as A) for whisking during epoch t1 when the wall is within whisker reach during CL. **C)** Same as A) for whisking during epoch t2 when whisker movement is restricted by nearby wall. **D)** Example 500ms segment of whisker angle (top) and whisker curvature (bottom) extracted from videos in A) for epoch t0. **E)** Same as D) for epoch t1. Middle: times of whisker contact with the wall. Large curvature increases correspond to whisker-wall touch events. **F)** Same as D) for epoch t2 when the wall is closest to the mouse. Amplitude is significantly reduced but curvature fluctuations remain. **G)** Set point comparison across the 3 epochs for n=10 randomly selected trials. **H)** Whisking amplitude comparison across the 3 epochs for the same 10 randomly selected trials, with a significant decrease seen during t2 (two sample *t*-test, p= 7.3×10^-5^). **I)** Comparison of whisker contact time with the wall for t1 and t2 epochs. When the wall is at its closest position, the whisker remains in contact for significantly longer durations (two sample *t*-test, *p* = 1.27×10^-4^). **J)** Comparison of whisking frequency for epoch t0, t1, and t2 (two sample *t*-test, t0 and t1 *p* = 0.259, t1 and t2 *p* = 0.147).

### Naturalistic posture generates phenotypic trotting gait at high-speed VR navigation

Rodents in the wild produce a specific set of gaits (walk, trot, gallop, and bound) that are expressed in distinct ranges of speed with phenotypic inter-limb and intra-limb coordination (Bellardita and Kiehn, 2015). Recruitment of distinct locomotor circuits at different speeds controls the alternation between left and right limbs. Switching between gaits occurs instantaneously as animals are navigating paths with varying slopes and frictions and adjusting body posture to safely redistribute body weight between the limbs. The light weight of the sphere in our VR (60g) and relatively small friction due to air-suspension (Methods) allow even mice with weight as small as 18g (n=26, 21.9g ± 3.03, mean ± s.d., **Table S1**) to easily turn it around during locomotion. To ensure the whisker-guided navigation in our rig is ethologically relevant, the position of the head-fixation bar was designed to be adjustable (**Fig.3A**) with head height (22mm) tuned to match head-free high-speed locomotion (Mitrevica and Murray, 2023). The sphere size (16-inches) is large enough to reduce the body tilt to a moderate 15° (**Fig.3A**) and thus reduce hunched postures typical of head-fixed locomotion along upward slopes in smaller size VRs(Dombeck et al., 2010). The resulting snout to hump angle (**Fig.3A**) closely matches 150° observed in head-free mice locomotion on a flat surface (Mitrevica and Murray, 2023) with nearly equal distribution of weight between fore and hind limbs.

**Figure 3.**
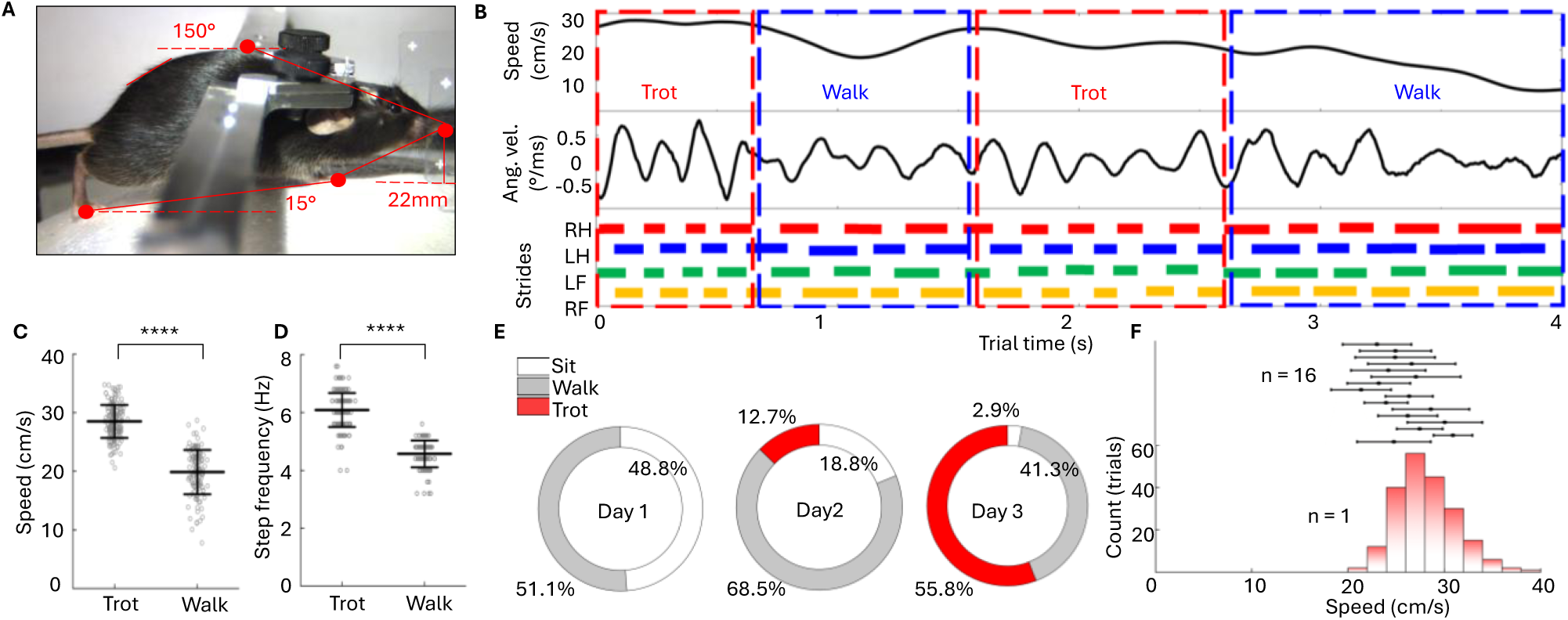
Naturalistic posture generates phenotypic trotting gait at high-speed during VR navigation. **A)** Mouse posture in VR rig matching the head height position and snout-to-hump angle of freely running mice. **B)** Example segment of straight CL single-whisker-guided navigation. Top: average mouse speed. Middle: angular velocity. Bottom: reconstruction of strides from videos with each leg’s contact times marked by colored lines. Red and blue dashed rectangles show instantaneous switching of gaits between trot and walk. **C)** Statistical comparison of mean speeds for trials classified as either walking or trotting. Two-sample *t*-test, *p* = 9.2E-68, n = 1, 200 trotting trials, 150 walking trials. **D)** Statistical comparison of stride frequency for trotting and walking, *p* = 5.8E-76, n = 1, 200 trotting trials, 150 walking trials **E)** Percentage of gaits used throughout habituation period for an example animal. **F)** Bottom: Histogram of speed for an example animal for trotting during main session. Top: Means and standard deviations trotting speeds for 16 animals.

To track animals’ gaits during navigation, three additional cameras illuminated with infrared LEDs were used to record positions of hind and fore legs (**Fig.3B**). Videos were synchronized and the time stamps when each paw was in contact with the sphere during strides (stance phase) were extracted manually (Methods) (**Fig.3B**, bottom). Individual strides can also be extracted (Sofroniew et al., 2014) from characteristic positive and negative peaks in angular velocity *dθ/dt* (**Fig.3B**, middle). We found mice switched between two main gaits, walk and trot, at fast speeds and occasionally many times during a single trial. Trotting was characterized by the diagonal limbs striding in synchrony, while walking was at slower speeds without such synchrony and with longer stance periods of contact between paws and the sphere. This difference is notably seen in the angular velocity trace, with trotting showing a periodic sinusoidal modulation (**Fig.3B**, middle), while walking corresponds to less synchronized lower amplitude modulation. To accurately quantify and classify gait as either trotting or walking, wavelet analysis was performed on the angular velocity trace, showing that lower frequencies (<4Hz) were present in only walking trials and absent during trotting (**Fig.S1C**). Stride frequency and mean speed were calculated for each trial and a Fisher’s linear discriminant analysis (LDA) was performed to classify segments as either walking or trotting (**Fig.S1B**). Trotting was performed at significantly faster speeds, 28.5cm/s ± 2.8 (mean ± s.d.), compared to walking trials at 19.9cm/s ± 3.8 (**Fig.3C**). Trotting was also carried out at faster step frequencies (6.1Hz ± 0.59, mean ± s.d.) compared to walking trials at 4.6Hz±0.46 (**Fig.3D**). These characteristics are consistent with gait speeds and stride frequencies observed during head-free locomotion (Bellardita and Kiehn, 2015).

We found mice preferentially perform single-whisker-guided navigation by trotting for increasingly longer times across habituation sessions starting from mostly walking in day 1, followed by trotting 12.7% time during session 2, and up to 55.8% time at the end of the habituation (**Fig.3E**). This was consistent across all mice (n=17), with final mean trotting speeds ranging from 23.8cm/s to 30.1cm/s (**Fig.3F**). Therefore, adjustments to the head-fixed posture in our VR rig resulted in generation of naturalistic gaits at high-speed. In what follows, only those trials during the main navigation session assigned as trotting, were used for analysis to eliminate inconsistencies attributed to differences in locomotion.

### ANC single-whisker-guided navigation-based decisions are executed within 250ms

Once animals are habituated, a perceptual navigation-based decision task is introduced to the paradigm (Methods) during a single and terminal main session that lasted up to 2 hours (**Fig.1C**). Each trial consists of two parts (**Fig.4A**, top): a section of straight CL wall tracking for 100cm forward distance as in the habituation sessions, followed by a 2 second-long open loop (OL) section when the CL feedback is terminated, and the walls are first briefly removed beyond the reach of the C2 whiskers and then just one of the walls rapidly approaches the snout (within 0.2s) and stays at a pre-defined constant distance for 1.8 seconds before it retracts again. Depending on which wall, right or left, is brought in closer to the animal, the mouse senses the approaching wall with its whiskers and is forced to turn either to the left or to the right, correspondingly, to avoid the expected collision, thus creating a version of a classical two-alternative forced choice (2AFC) task (**Fig.4A**, bottom, see also **Movie S1**). However, typical 2AFC paradigms require prolonged weeks-long training and introduction of rewards or punishments for learning enforcement. In contrast, our naturalistic task is aimed to mimic closely the natural navigation mice perform in burrows, when animal navigates left and right turns of the burrow without expecting rewards. Therefore, while we use water rewards in the habituation sessions (Fig.1C) to encourage running, we intentionally blocked them in the main behavioral session when the animal is introduced to turns. Therefore, we refer our task as an Alternate Navigation Choice paradigm (ANC), where the outcomes are to successfully execute corresponding turn thus keeping the binary decision framework of classical AFC.

**Figure 4.**
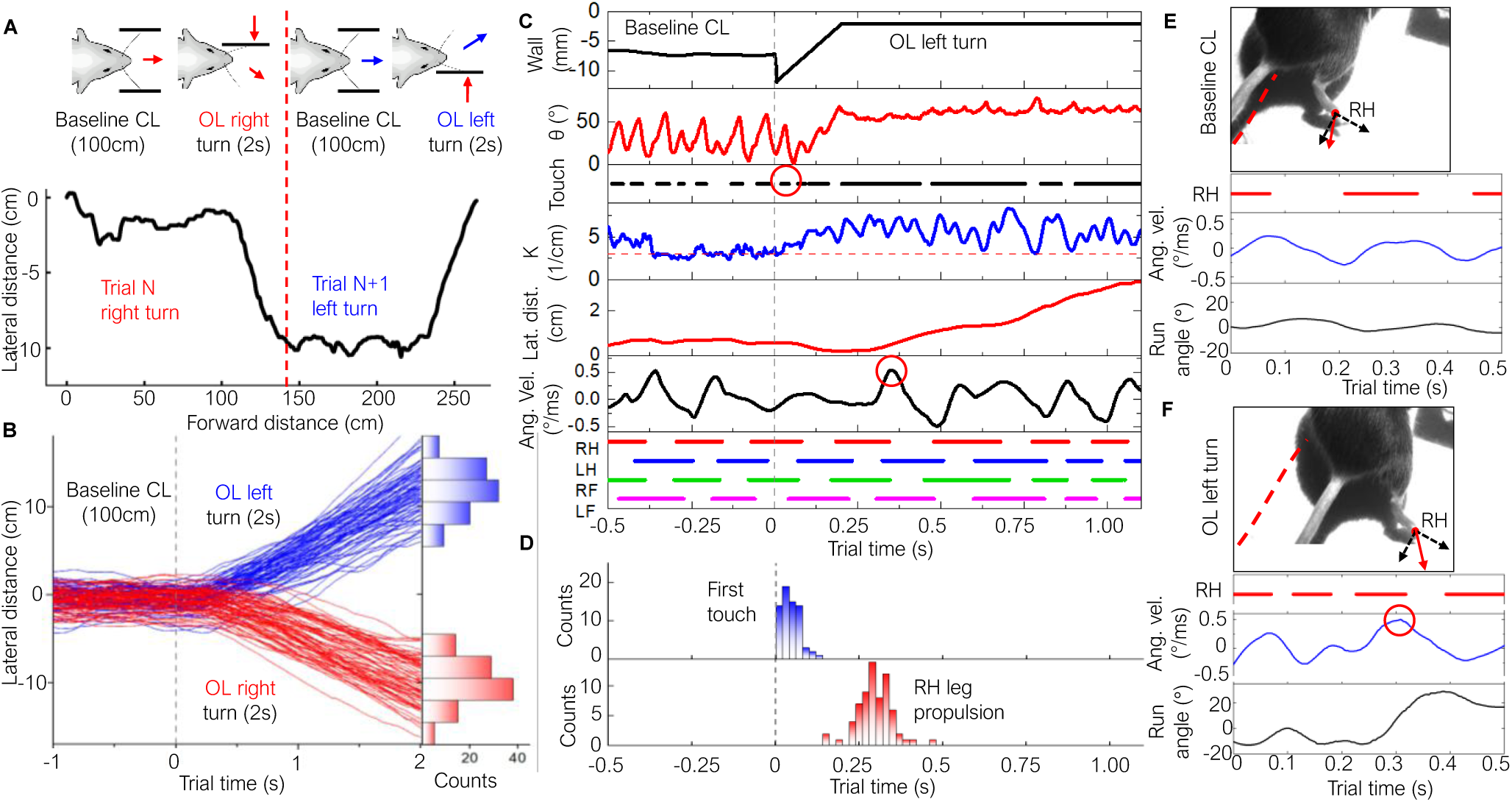
ANC single-whisker-guided navigation-based decisions are executed within 250ms. **A)** Top: Schematics of the trial structure for a left and right turn, showing 100cm straight running in closed-loop environment before walls are briefly retracted and then one wall is moved closer to the mouse for 2 seconds forcing mouse to turn in response. Bottom: Example mouse trajectory recorded during two consecutive trials with OL Left and OL Right turns. **B)** Overlaid trajectories of 143 randomized OL Left (blue) and OL Right (red) trials during continuous 2-hour unrewarded navigation. Right: histogram showing resulting lateral distance distribution for both trial types. **C)** Tracked behavioral parameters during example left turn decision. Wall position (black, top) shows right wall approaching the mouse. Whisker angle (red), wall touch events (black), and curvature (blue) are extracted from video. The red circle corresponds to the first whisker touch to approaching wall. Lateral mouse distance (red) and angular velocity (black) are extracted from tracking the sphere positions. The red circle corresponds to RH leg propulsion. Bottom: Stance strides extracted from simultaneous tracking of all 4 paws. **D)** Distributions of decision onset (first whisker touch to the approaching wall) and decision execution (maximum propulsion during the turn) for the same example mouse. **E)** Top: Videoframe showing position of RH leg during propulsion at the end of a stance during CL straight run. The red dashed line is forward direction. Bottom: Example segment showing relationship between the RH leg stance (red), angular velocity (blue), and run angle (black). **F)** Same as E) during OL Left turn. The red circle corresponds to RH leg propulsion at the end of the stance.

Typical trajectories of an example animal navigating left and right turns during 2-hour long session (**Fig.4B**, left, 143 trials) with overlaid sections of straight runs (CL, negative times) and randomized left (OL Left, blue curves, positive times) and right (OL Right, red curves, positive times) turns are all performed while the animal is running at elevated speed. While during the CL straight section animal is staying within ±2cm success criterion (1.2cm s.d.), trajectories during turns end up at 10.2cm ± 2.9 and -10.2cm ± 2.6 (mean ± s.d.) for left and right turns, correspondingly (**Fig.4B**, right). A turn success criterion can be applied by quantifying whether the mouse has turned by a minimum of 10° change in run angle at the end of the trial compared to the CL section. For the session on **Fig.4B** the resulting success rate is 82%.

To identify critical times of decision-making when the animal is receiving the first information that the wall is fast approaching up to the time when a decision to turn is executed, we analyzed synchronized recordings of paw positions and whiskers touch events (**Fig.4C**). The first touch (FT) of a whisker to the approaching wall (**Fig.4C**, red circle) can be detected at 30ms after the start of the turn section that is followed by an increase of curvature amplitude characteristic of the switching to epoch t2 of whisking restricted by a close-by wall (**Fig.2C**). Histogram of FT moments (**Fig.4D**, top) reveals narrow FT distribution with 46.0ms ± 27.4 (mean ± s.d.). Note, that whisker is constantly palpating the wall during the CL section (epoch t1) prior to the FT. Therefore, unlike many 2AFC paradigms, the FT manifests the arrival of a deviant odd-ball change in the otherwise continuous sensory signal, rather than a sudden onset of previously absent stimulus.

The moment of motor execution of a decision to turn (reaction time, RT) is detected as a characteristic peak in the angular velocity *dθ/dt* (**Fig.4C**, bottom, red circle). Gait reconstruction during individual strides (**Fig.4C**, bottom, colored rectangles) aligns the RT to the end of a stance phase in a diagonal trot. At this time the body axis (lateral displacement of base of tail) that was initially aligned with the straight run direction (**Fig.4E**, red dashed line) is changed to provide the largest propulsion (**Fig.4F**) by the right hind (RH) leg in the 35° direction of the turn. The maximum angular velocity corresponds to the end of the stance phase (full stance to toe push off) when the RH leg is leaving the ground with strongest propulsion (**Fig.4F**, bottom) along the direction of the turn. Such stereotypic thrust is observed for all left and right trials (**Fig.S2A**). For the experiment of **Fig.4B** the RT is 295.1ms ± 52.9 (mean ± s.d.) as shown in the RT histogram in **Fig.4D**, bottom. Therefore, the total duration of the decision period from the onset (FT) to the execution (RT) can be estimated for this example animal as 249.1ms ± 52.9 (mean ± s.d.). Note that animals were not previously exposed to this task. However, they start to execute turns correctly from the first time the turns are encountered (**Fig.S2B**). The task is also not rewarded as the water-spout is moved out of reach. Therefore, our ANC task is neither learned nor rewarded.

### Calibrated lesioning of small subpopulation of neurons in layer 4 of a C2 cortical barrel column

Previous studies have shown that the barrel cortex is dispensable for some whisker guided behaviors including vibration discrimination (Hutson and Masterton, 1986), passive touch detection(Hong et al., 2018; Waiblinger et al., 2022), and detecting obstacles (Warren et al., 2021), but required for other tasks including aperture discrimination (Krupa et al., 2001), gap crossing(Hutson and Masterton, 1986; Shih et al., 2013), texture discrimination (Guic-Robles et al., 1992), and whisker-guided navigation (Sofroniew et al., 2015) tasks.

To identify the contribution of wS1 circuits to decisions in our single-whisker-guided VR navigation paradigm, we developed a localized electrolytic lesioning technique (Methods). To produce lesions localized within the principal C2 barrel column, we utilized a multielectrode array (MEA) probe aligned perpendicular to the cortical layers. To avoid stereotaxic surgery and enable targeted probe insertion and lesioning during behavioral experiment we used *Scnn1a-TG3-Cre* x *Ai32* transgenic mice. Strong fluorescence from yellow fluorescent protein (EYFP) expressed in layer 4 spiny stellate cells(Madisen et al., 2012) enables observation of the entire wS1 *in-vivo* to target the principal C2 whisker barrel (**Fig.5A**). Once the probe is inserted in the left hemisphere aligned parallel to the C2 barrel column (**Fig.5B-D**), optogenetic tagging is performed by exciting co-expressed Channelrhodopsin-2 (ChR2) to obtain a current source density (CSD) plot (**Fig.5E**) in which a sink originates in the center of layer 4. We then retract the probe to align its last electrode to the center of layer 4 and perform lesion by passing the calibrated current (Methods) to the positioned last electrode. As opposed to previous studies when large volumes of barrel cortex were inactivated with potential confounding off-target effects (Hutson and Masterton, 1986; Krupa et al., 2001; O’Connor et al., 2010; Sofroniew et al., 2015; Chakrabarti and Schwarz, 2018; Hong et al., 2018; Warren et al., 2021; Ryan et al., 2022; Waiblinger et al., 2022; Gardères et al., 2024), we calibrated the lesioning to target affected cortical volumes of 10^-3^mm^3^ (**Fig.S3A**). *Ex-vivo* histological mapping of the whole brain (**Fig.5B**) and of coronal slices (**Fig.5D**) confirms the probe placement and lesion position. The lesion and barrel volumes as well as cell counts were quantified via confocal microscopy using histological staining (**Fig.5F-H, Fig.S4A-B**) that enables volumetric reconstruction of the barrel and the embedded lesion (**Fig.5I**). For reference, similar reconstruction was done on the intact C2 principal barrel column in the right hemisphere (**Fig.S4C**) with the estimated volume of L4 barrel of 9×10^-3^mm^3^. Cell counting (**Fig.5H, Fig.S4C**) results in 1250cells ±100 (mean ± s.d.) that corresponds to cell density of 1.4×10^5^cells/mm^3^.The estimated volume of a lesion (**Fig.5I**) of 9×10^-4^mm^3^ (Methods) constitutes therefore about 10% of the L4 barrel volume that provides an estimate for the affected neuron soma just above 100. Our cell density numbers are consistent with numbers reported previously (1800 cells(Lefort et al., 2009) with densities (Herculano-Houzel et al., 2013; Keller et al., 2018) between 1.3×10^5^cells/mm^3^ and 2×10^5^cells/mm^3^) that provides an upper limit of affected neuron soma below 200 It is important to note that besides damaged cells, the volumetric lesion also disrupts afferent and efferent columnar connections (neuron axons and dendrites) as well as non-neuronal cells (microglia, astrocytes, etc.) that may also contribute to the loss-of-function. In addition, lesion can disrupt microvasculature that was shown to degrade whisker-guided gap crossing behavior with disruption of a just single penetrating vessel (microinfarctions volume of 28% of the cortical column) (Shih et al., 2013).

**Figure 5.**
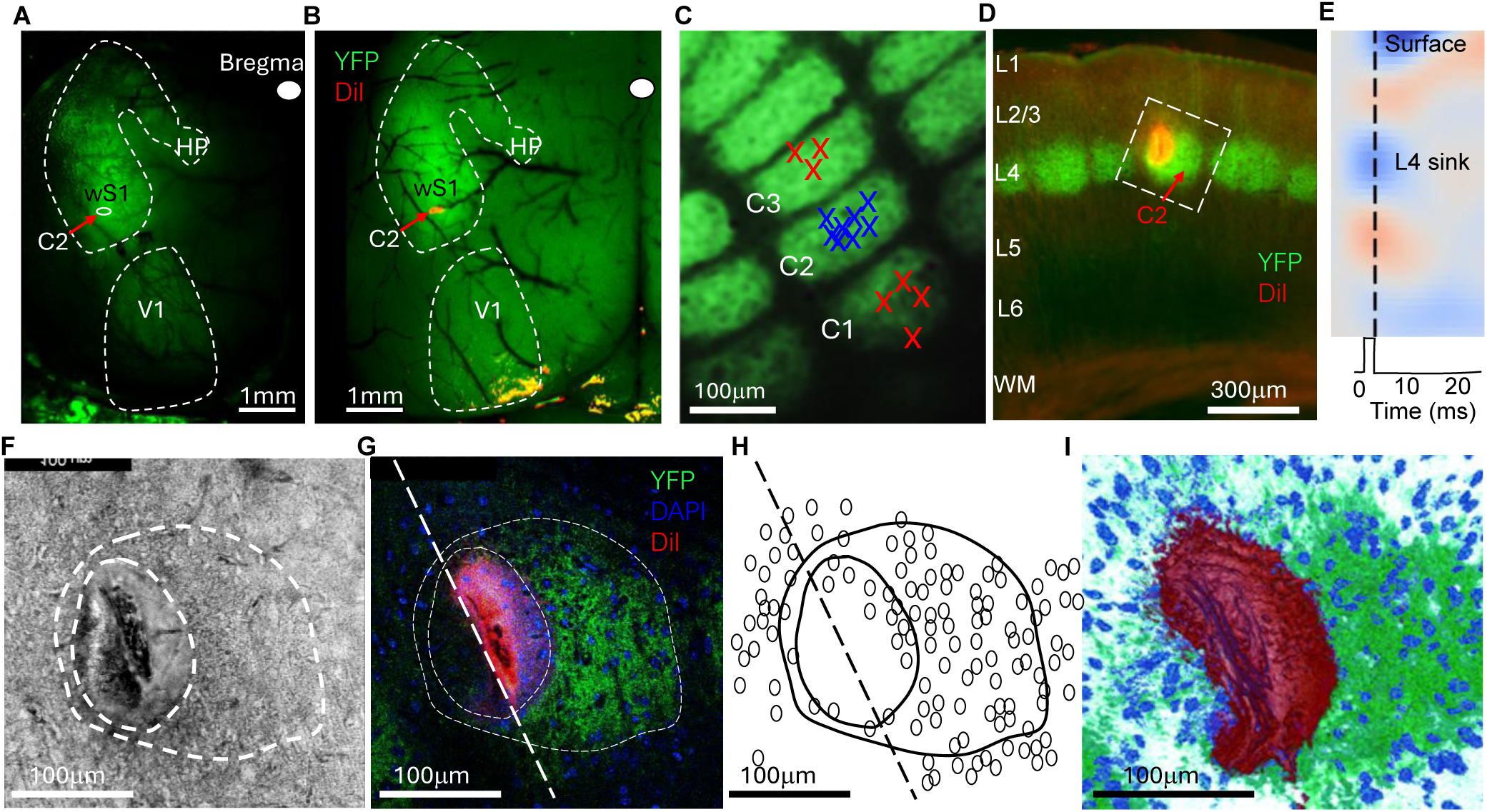
Targeted insertions and subsequent lesion quantification. **(a)** Top down in-vivo fluorescent microscope image of the animal skull though the open headpost window showing boundaries of V1, and somatosensory wS1 and HP areas. **(b)** Top-down fluorescent microscope image of the partially perfused and fixed brain. Probe track is marked by DiI red trace to visualize the entry point. **(c)** Tangential section of the barrel cortex with overlaid reconstructed implantation sites for principal barrel C2 and neighboring C1 and C3 barrels. **(d)** Overlaid fluorescence microscope images of the coronal 50µm thick section, for EYFP (green) and DiI (red) filters showing location of cortical layer’ boundaries, C2 barrel, and electrolytic lesion. Whie rectangle shows the orientation of the image in (f). **(e)** Current Source Density map produced during *in-vivo* opto-tagging to identify the location of L4 sink for lesion targeting. **(f)** Magnified bright field image of a lesion image in (d) taken with confocal microscope, with lesion and layer 4 of the C2 barrel column outlined with white dashed line. **(g)** Overlaid confocal microscope images of the same slice as in (f). A single coronal section from a Z-stack is shown with a C2 barrel (YFP, green), decorated lesion (DiI, red), and cell nuclei (DAPI, blue). Dashed white line is a probe track. **(h)** Map of C2 L4 barrel boundaries, lesion size, and cells ROI for the same slice extracted from (g). **(i)** Volumetric reconstruction of a lesion within L4 principal C2 barrel extracted from a full confocal Z-stack using 3D slicer.

### Local lesion in layer 4 disrupts single-whisker-guided navigation during contralateral but not ipsilateral turns

To test the loss-of-function consequences of the localized lesion on navigation decisions, the ANC protocol was implemented during the main session (Methods). Once habituated, animals are head-fixed on the VR rig and start to run freely with walls out of whiskers reach, the MEA probe was inserted in either the center of a C2 barrel column or a neighboring barrel columns following previously acquired barrel field map (**Fig.5C**). Since the probe is always inserted into the left hemisphere, the left and right turn trials can be identified as contralateral and ipsilateral,

To provide a baseline for loss-of-function analysis of behavior, we performed experiments with a cohort of animals (n=7) in which a L4 lesion was produced not in the principal C2, but in the neighboring C1 and C3 barrels (**Fig.6A-D**), which are located within 100μm distance in the barrel cortex (Fig.5C) and belong to the same C-row of whiskers oriented parallel to the ground. The experiment consists of 2 consecutive sessions running back-to-back with under 2-minute interval. The first session is to acquire baseline behavior during a 10-minute ANC session with whisker-guided left and right turn decisions (**Fig.5A**). This is followed by a calibrated charge (**Fig.S3A**) applied to the lesion electrode, while animal is actively navigating with the whole procedure lasting less than 75 sec. No noticeable behavioral changes were observed during the procedure. Immediately afterwards, a post-lesion 10-minute ANC session was carried out with the same left and right turns (**Fig.5B**). Comparison of performance for contralateral left turns during session 1 before the lesion (**Fig.5A**, blue) showed success rate of 82.3%±9.2 (mean ± s.d.) and 72.7%±7.6 (mean ± s.d.) after the lesion (**Fig.5B**, blue). For ipsilateral right turns (**Fig.5A**, red) success of 69.4%±10.9 (mean ± s.d.) before lesion is statistically similar to 72.9%±18.2 (mean ± s.d.) acquired after the lesion is made (**Fig.5B**, red). This is consistent across all animals in this cohort (n=7) with no significant difference (p>0.05, paired sample t-test) observed for both left contralateral (**Fig.5C**) and right ipsilateral (**Fig.5D**) turns.

**Figure 6.**
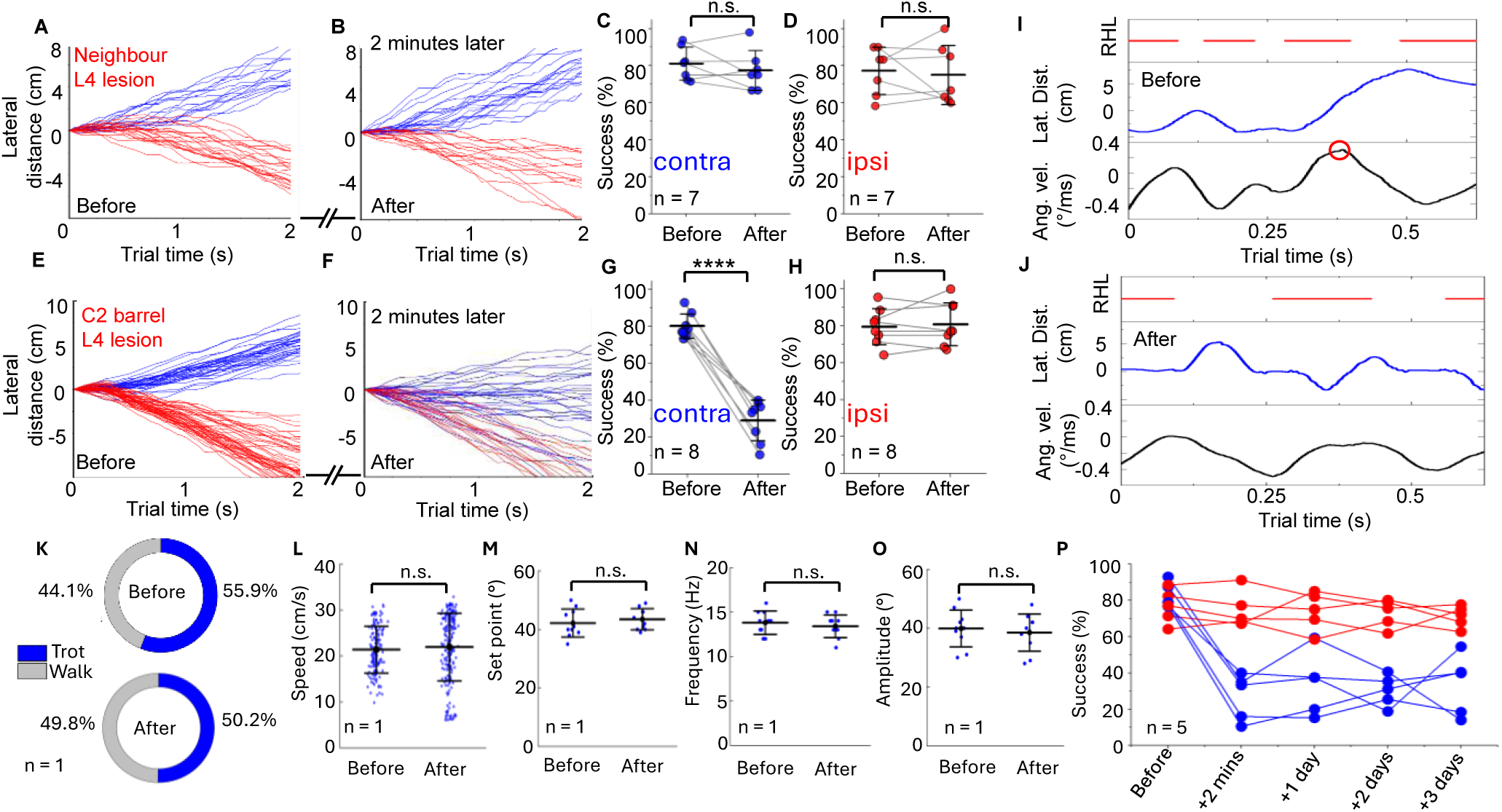
Single-whisker-guided navigation decisions causally depend on small subpopulation of neurons in layer 4 of the principal barrel column. **A)** Overlaid trajectories for left (blue) and right (red) turning trials for a ten-minute long baseline behavioral session 1 for an example animal. **B)** Ten-minute long behavioral session 2 for the same animal recorded 2 minutes after the lesion is produced. The lesion was located not in the principal, but in the neighboring C1 barrel. **C)** Success rate in contralateral (left turns) trials of 7 animals before and after lesions in neighboring barrels showing no significant difference, pair sample *t*-test, *p* = 0.58. **D)** Success rate in ipsilateral (right turns) trials of 7 animals before and after lesions in neighboring barrels showing no significant difference, pair sample *t*-test, *p* = 0.12. **E)** Ten-minute long baseline behavioral session 1 for another example animal with the trajectories for left (blue) and right (red) turning trials overlaid. **F)** Ten-minute long behavioral session 2 for the same animal recorded 2 minutes after the lesion is produced. The lesion was located in the principal C2 barrel. **G)** Success rate in contralateral (left turns) trials of 8 animals before and after lesions in the principal C2 barrel showing significant decrease in behavioral performance, pair sample *t*-test, *p* = 0.006. **H)** Success rate in ipsilateral (right turns) trials of the same 8 animals before and after lesions in the principal C2 barrel showing no significant difference, pair sample *t*-test, *p* = 0.95. **I)** Top: RH leg stance (red); Middle: Lateral distance (blue); Bottom: angular velocity (black) during the decision period for a contralateral (left) trial in an example animal before a lesion in the C2 barrel column, showing characteristic angular velocity peak (red circle) and subsequent left turn in response to the approaching wall. **J)** Same plots for the same animal after a lesion in the C2 barrel column is made. Animal ignores closely approaching wall while continuing undisrupted locomotion. **K)** Percentage of behavioral session spent walking or trotting before and after lesion in an example mouse. **L)** Comparison of trial-by-trial mean speeds before and after lesion in C2 barrel layer 4 for a single animal, two sample *t*-test, *p* = 0.42. **M)** Comparison of mean whisking set point for 10 randomly selected trials before and after C2 layer 4 lesion for the same single animal, two sample *t*-test, *p* = 0.50. **N)** Comparison of mean whisking frequency for 10 randomly selected trials before and after C2 layer 4 lesion for the same single animal, two sample *t*-test, *p* = 0.50. **O)** Comparison of mean whisking amplitude for 10 randomly selected trials before and after C2 layer 4 lesion for the same single animal, two sample *t*-test, *p* = 0.63. **P)** Recovery tests, showing success rates in contralateral (blue) and ipsilateral (red) trials for 5 animals in the same behavioral sessions spanning 3 days post-lesion.

In striking contrast, for a cohort of animals (n=8) with the L4 lesion produced in their C2 principal barrel column that corresponds to the only pair of C2 whiskers they are navigating the VR with, we observed immediate degradation of performance. For contralateral left turns the success rate of 83.4%±8.1 (mean ± s.d.) acquired during session 1 before the lesion (**Fig.5E**, blue), drops dramatically down to 31.7%±7.2 (mean ± s.d.) in session 2 after the lesion (**Fig.5F**, blue). Loss of performance (**Fig.5G**) is statistically significant (paired *t*-test, *p* = 0.006) across all animals in this cohort. However, quite remarkably, the performance in the ipsilateral turns remained intact with the baseline success rate at 81.3%± 10.7 (mean ± s.d.) (**Fig.5E**, red) before the lesion remaining at 80.9%±18.7 (mean ± s.d.) (**Fig.5F**, red) after the lesion. Across all animals in this cohort (**Fig.5H**), it remains statistically indistinguishable (two-sampled t-test, p>0.05).

### Single-whisker-guided navigation causally depends on sensory information flowing through L4 of a single principal whisker barrel

At this stage it is important to identify possible off-target effects of the lesion that may affect behavioral performance and confound the interpretation. First, we explored whether cortical lesions may affect the execution of decisions during locomotion. Tracking paws positions during decision making in session 1 prior to the lesion shows characteristic peak in angular velocity at the end of a stance phase (**Fig.5I**, red oval) associated with strong propulsion delivered by the RH leg. In the example trial directly following the L4 lesion in principal C2 barrel, the animal continues to run along the previous direction (**Fig.5J**, blue) executing regular trot strides (**Fig.5J**, black) practically ignoring the wall that approached the snout to a distance of just 2mm. Major locomotion parameters such as percentage of the session spent walking or trotting (**Fig.5K**) and the mean speed across trials (**Fig.5L**) also remain intact with no significant difference (two-sampled t-test, p>0.05).

Second, we explored whether the L4 lesion affects the whisking motor action, as barrel wS1 cortex is known to be involved in regulation of active whisking(Kleinfeld et al., 2006; Feldmeyer et al., 2013; Staiger and Petersen, 2020; Diamond and Toso, 2023). For that, 10 trials were randomly selected to extract set point (**Fig.5M**), frequency (**Fig.5N**), and amplitude (**Fig.5O**) from overhead videos from session 1 prior to lesion and session 2 immediately following lesion. In all cases no statistical difference was detected (paired t-test, p>0.05), indicating that whisking motor action is not affected.

Next, we explored whether electrical stimulation during lesion formation can by itself suppress cortical activity thus potentially contributing to observed degradation of behavior. Strong cortical inhibition lasting 200-400ms has indeed been frequently observed following brief sub-millisecond electrical stimulation (Berman et al., 1991; Chung and Ferster, 1998). In our case of a few seconds long stimulation, it is unclear whether the cortex will remain functional minutes after the lesion is formed. To address this, we performed electrophysiological recordings of spiking activity (Methods) spanning the whole principal C2 barrel column (n = 7) for a 20 min session before the lesion and compare them to recordings performed in a 20 min session immediately after the lesion is formed (**Fig.S3C-E**). For a single animal, the neural spiking activity measured across the whole column indeed exhibits substantial suppression immediately following the lesion (**Fig.S3C**). However, this inhibition is short lived with a complete recovery to the pre-lesion values in just a few minutes. Outside of this transient period, the neural activity before the lesion (9.3 ± 0.1 sp/sec/unit, mean ± SEM) and following 2 min after the lesion (9.1±0.1) is statistically identical (**Fig.S3D**), which is consistent across all n=7 mice (**Fig.S3E**). Despite the return of spiking activity to pre-lesion values, however, the behavior remains disrupted (**Fig.6G, Fig.S3B**). Therefore, we conclude that off-target effects of the lesion procedure itself do not contribute to the observed changes in behavior.

Lastly, we explored whether the loss-of-function can be recovered after adaptation and learning. Previously rapid recovery was observed after inactivation of barrel cortex(Hong et al., 2018; Warren et al., 2021) that indicates temporal disruption of sensory flows between areas primarily involved in the decision making followed by experience-dependent reconfiguration. In contrast, in our paradigm we found that sudden drop of success rate for contralateral left turns immediately following a lesion in the principal C2 barrel column from 93.0% to 34.9% (**Fig.5P**) stayed low at 37.4±18.3 (mean ± s.d.) for this cohort of mice (n=5). This indicates that the most plausible scenario for loss-of-function is not the temporal disruption of sensory flows to other brain areas, but the inability to access the salient sensory information itself to make a decision.

What will happen if, by increasing the lesion volume, larger populations will be affected? To explore this have carried out additional experiments with a range of lesion volumes produced by different doses (**Fig.S3A**) and assessed their impact on behavioral performance (**Fig.S3B**). Even a small lesion of less than 10% of the L4 barrel volume results in significant degradation of success rate below 40%. Interestingly, increase of the lesion volume up to 50% shows no statistically significant changes. Further increase of the lesion volume approaching 80-90% of the L4 barrel significantly degrades the performance down to complete loss of the ability to navigate. We speculate that this observation is related to various effects of a volumetric lesion on disruption of neuron processes degrading communication channels throughout the barrel column, an increasing number of affected neuron soma, involvement of other non-neuronal cells, and microvasculature damage, probably in this order. Note, that even the largest lesion volumes studied in this work that cover the whole L4 barrel are much smaller than reported in the literature including the closest report on a single penetrating vessel microinfarction (Shih et al., 2013) and its effect on a similar ethological behavioral task of whisker-guided gap crossing (28% of the whole cortical column volume).

All of these observations considered jointly enables us to conclude that single-whisker-guided navigation in our VR causally depends on activity within a small neuronal subpopulation in a L4 of a single principal C2 barrel. Therefore, by designing our whisker-guided navigation paradigm and restricting sensory inputs to just a single whisker, we effectively generated an information bottleneck, that, when destroyed, could not be recovered by redistribution of the information flows to other brain areas.

## Discussion

We have shown that the principal barrel column is necessary for single whisker-guided navigation-based decisions, with successful navigation relying on small subpopulation of neurons in the input layer 4. Why then does this behavioral task require the barrel cortex, whereas other whisker-dependent behaviors do not?

Animal navigating the turns is receiving information from its whiskers on fast approaching wall (**Fig.7A**). An internal model of the body position (Wolpert et al., 1995) is constantly updated via proprioception. A single FT event informs the animal to predict moments in time when different body parts (**Fig.7A**) will be in a colliding path with the approaching obstacle – so called two-dimensional time-to-collision (2D TTC) used in a simple forward control of autonomous driving vehicles(Hayward, 1971) (ADV). Calculated 2D TTC (**Fig.7B**) gives an animal about 300ms for clearance of the RH paw and just 50ms to clear the RF paw as it is closer to the wall. We argue that many whisker-dependent behavioral paradigms fall into this class of problems including detection of whisker touch(Hong et al., 2018; Waiblinger et al., 2022), vibration discrimination (Hutson and Masterton, 1986), and evading obstacles that were reported by a single whisker touch event (Warren et al., 2021). The 2D TTC is a relatively simple forward control algorithm that relies on internal models of body position but not necessarily associated with the neural activity in wS1. While collision prediction requires continuous updates of the internal body model, it does not however require updates of the obstacle position by whisking therefore making the wS1 potentially dispensable.

**Figure 7.**
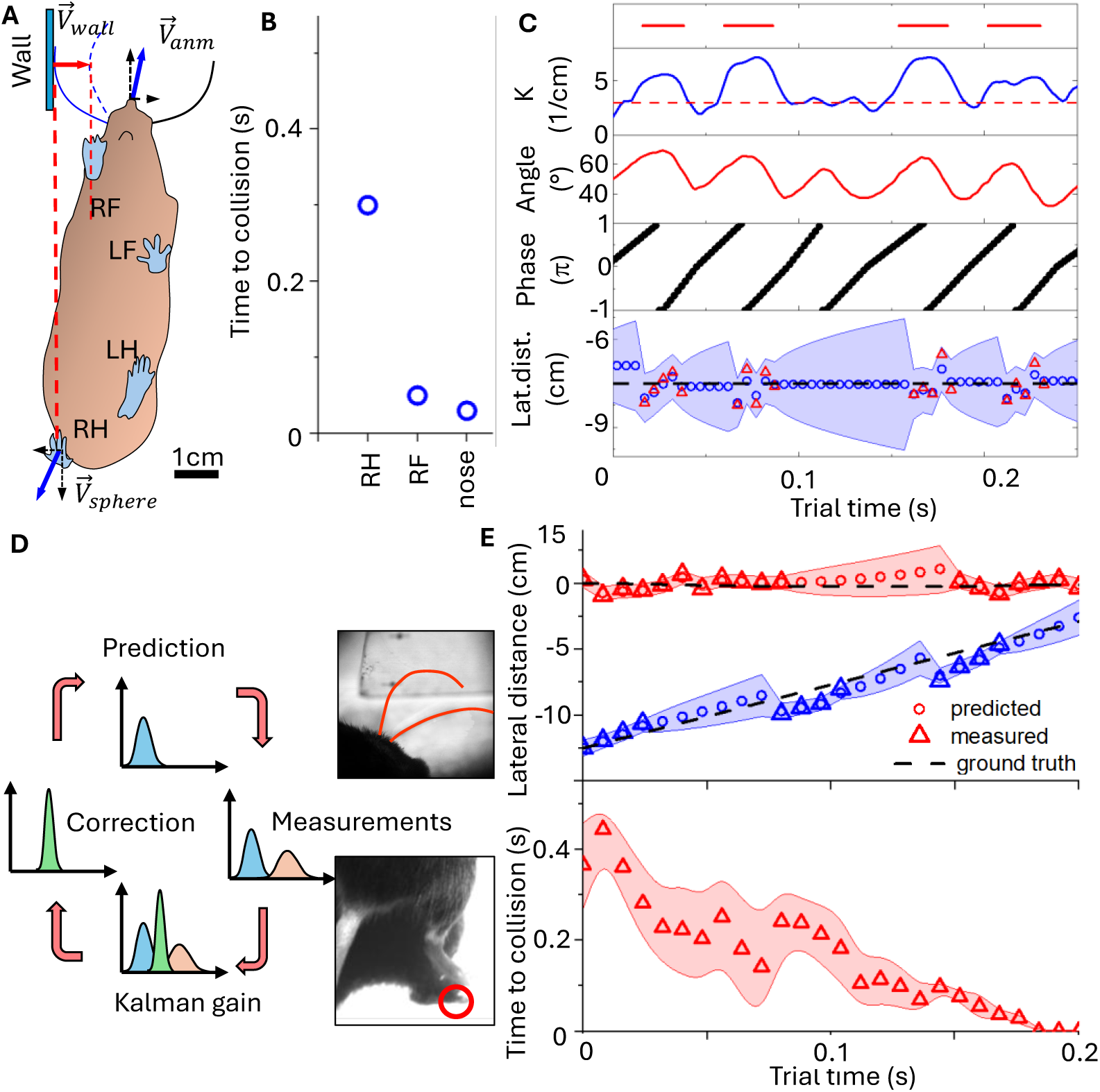
Probabilistic navigation in the presence of noise and occlusion of measurements. **A)** Schematics (view from below animal) of a mouse making the decision to turn when whisker senses the approaching wall. **B)** Deterministic 2D TTC calculated for schematics in A) for RH leg, RF leg, and for nose colliding with the approaching wall. **C)** Example of a CL session with one touch event (t=0.12s) skipped. Top: Whisker contacts the wall. Middle: Whisker curvature (blue), angle (red), and whisking phase (black). Bottom: Kalman filter predictions of the wall position (blue circles) compared to actual GT trajectory (dashed black), and measured positions (red triangles). Shaded region corresponds to 1 s.d. gaussian distribution. **D)** Schematics of discrete Kalman filter. Measurements of whisker curvature provide the wall state vector and its statistical distribution (pink gaussian distribution), while proprioception of leg positions provide corresponding state vector of the animal body. Internal models of body and wall movements enable prediction of their relative positions with corresponding uncertainties (blue gaussian curve) at the next time step and evaluate probability of collision with corresponding distribution. **E)** Probabilistic 2D TTC using Kalman filter. Top: Probabilistic predictions of positions of the wall (blue) and the animal body (red) for a section of OL left turn. Triangles correspond to measurements of the right wall positions when whisker is in contact (red) and to body positions when paws are in contact (blue). Circles represent predictions from Kalman filters that can be made even when measurements are not available. Shaded regions correspond to 95% confidence intervals. Predicted wall and body positions are close to the actual ground truth (black dashed lines). Bottom: Probabilistic 2D TTC. Shaded region corresponds to 1 s.d. gaussian distribution.

In contrast, the problem of tracking the walls and navigating the turns requires continuous updates on the states (position, speed, acceleration) of both, the body and the obstacle, with corresponding adjustment of the motor action following the change in their relative positions. ADV navigation in such environment is becoming a non-trivial task for which the forward control is insufficient, and a feedback control is required. The process should be tolerant of a large amount of noise introduced during measurements and should be robust even when measurements are not available. An example of such noisy and incomplete measurements is presented in **Fig.7C** for a section of a CL wall tracking trial. The wall distance can be decoded (Cheung et al., 2019) by the animal from whisker curvature (**Fig.7C**, blue) that induces stress-dependent spiking in primary afferents (Severson et al., 2017) followed by feature extraction in wS1 (Sofroniew et al., 2015). Such decoding (Kleinfeld et al., 2006; Feldmeyer et al., 2013; Staiger and Petersen, 2020) requires knowledge of whisking amplitude (O’Connor et al., 2010) (**Fig.7C**, red), phase(Isett and Feldman, 2020) (**Fig.7C**, black), set point, or all of them together to disentangle distance-dependent stresses from self-motion induced nonlinearities and is inevitably noisy. Moreover, the information on the wall state is available only when the whisker is in contact with the wall (just 40% of the time in a foveal mode, **Fig.2I**) and sometimes is lost when the touch event is missing (note absence of touch at 0.12s in **Fig.6C**) that leads to omission of whisker curvature change).

There is a strong consensus that brain performs probabilistic inference (Pouget et al., 2013) based on representations of probability distributions. Navigation based on sensorimotor integration in the presence of noise and uncertainty requires internal probabilistic models (Wolpert et al., 1995) of the dynamics of the body and external obstacles to predict possible collisions and produce motor actions to avoid them. It has been argued (Wolpert et al., 1995; Denève et al., 2007) that neural circuits may implement a close approximation to a Kalman Filter (KF) that is widely used in ADV feedback control systems (**Fig.7D**). To illustrate such probabilistic whisker-guided navigation we introduced (Methods) a discrete Kalman filter (KF) to model probabilistic predictions on wall positions (**Fig.7C**, bottom). We assume model linearity and gaussian noise in decoding of the wall state from whisker variables. The measurements are present (red triangles) only when whiskers are in contact with the wall and are updated each 8ms that roughly mimic the shortest duration of cortical feedback loops(Staiger and Petersen, 2020). The KF model produces (**Fig.7C**, bottom) reasonable predictions (blue circles) close to ground truth (GT) trajectory (black dashed line), however with increasing uncertainty (blue shades) when touch events are skipped. To emulate the internal decision feedback loop during whisker-guided navigation we used a 0.3s section from a ANC OL left turn experiment with recorded whisker variables and navigation parameters (**Fig.S5**) and known GT positions of the wall and a sphere (**Fig.7E**, dashed black curves). An animal is trotting during this section with RH leg in contact with the sphere every 140ms (**Fig.7E**, top blue triangles). For this trial the whisker first touches the approaching wall at FT of 7ms that is then repeated 3 times every 60ms before the decision to turn is implemented by the RH propulsion at the end of a stance.

We assume that measurements of the wall and body parts positions are accompanied by gaussian noise and are available only during their contact times sampled at KF cycle frequency. The resulting predictions (**Fig.7E**, red and blue circles) follow closely the GT trajectories. While probabilistic 2D TTC calculated using the KF predictions (**Fig.7E**, bottom) at the FT event (0.37s ± 0.09, mean ± s.d.) is close to the deterministic value (0.32s from **Fig.7B**), the KF produces continuous predictions even when the measurements of wall and body positions are out of sync or are not present. Extending analogy to ADV controls, computational resources needed to implement a probabilistic feedback control to navigate in the presence of uncertainty, noise, and occluded measurements, are typically significantly more demanding than for a simple forward control triggered by a single event. In fact, as of late 2024 no ADV system has achieved fully autonomous driving capabilities(Sever and Contissa, 2024) (level 5 of the SAE J3016 standard).

A possible alternative interpretation to consider is that the barrel cortex becomes engaged only to maintain normal behavior in a new and challenging environment, such as a head-fixed behavioral paradigm. However, even in head-free perceptual decision-making experiments (Hutson and Masterton, 1986; Shih et al., 2013) strong degradation of whisker-guided gap crossing is reported after ablation of the contralateral barrel cortex. Moreover, most prior loss-of-function experiments, either head-free or head-fixed, rely on weeks-long training before psychometric testing is performed. Therefore, at this late stage, the role of the cortex in learning the new task and the new environment has already been completed. In contrast, our “naturalistic” ANC paradigm requires a short 3-day habituation period (**Fig.1**) to produce natural running gait at high speed. The task itself – to produce a turn once the wall is sensed – is executed from the very first presentation of the turns (**Fig.S2B**) with high fidelity and negligible error rate that remains very low for the whole duration of the session. All our results, therefore, favor the hypothesis that barrel cortex plays a key role in generating probabilistic predictions as illustrated by our KF 2D TTC model (**Fig.7C,D,E**).

These findings may reconcile seemingly contradictory observations on the role of primary somatosensory cortex in perceptual decision making. Relatively simple forward control schemes triggered by a single sensory event to predict and avoid collision(Hong et al., 2018; Warren et al., 2021), may not require wS1 cortex as whisker-related sensory signals can report appearance of an obstacle using other information pathways. In contrast, whisker-guided navigation of turns in our VR paradigm requires continuous updates and predictions of relative positions of the body and obstacles in the presence of noise and missed measurements to infer the optimal path for collision avoidance. These complex computations are likely to rely on nested feedback loops that directly involve wS1 (Kleinfeld et al., 2006; Feldmeyer et al., 2013; Armstrong and Vlasov, 2025), especially when the sensory information is forced to flow through an information bottleneck in L4 of principal whisker barrel defined in our experiments by a single-whisker-guided paradigm.

## Acknowlegements

YV is a CZ Biohub Investigator. Authors are grateful to Kun Hu and Nur Al-Kodmany for help in setting up the VR rig and development of behavioral protocol and to Alexander Lee for help with KF coding.

## Supplemental Information

**Table S1.**
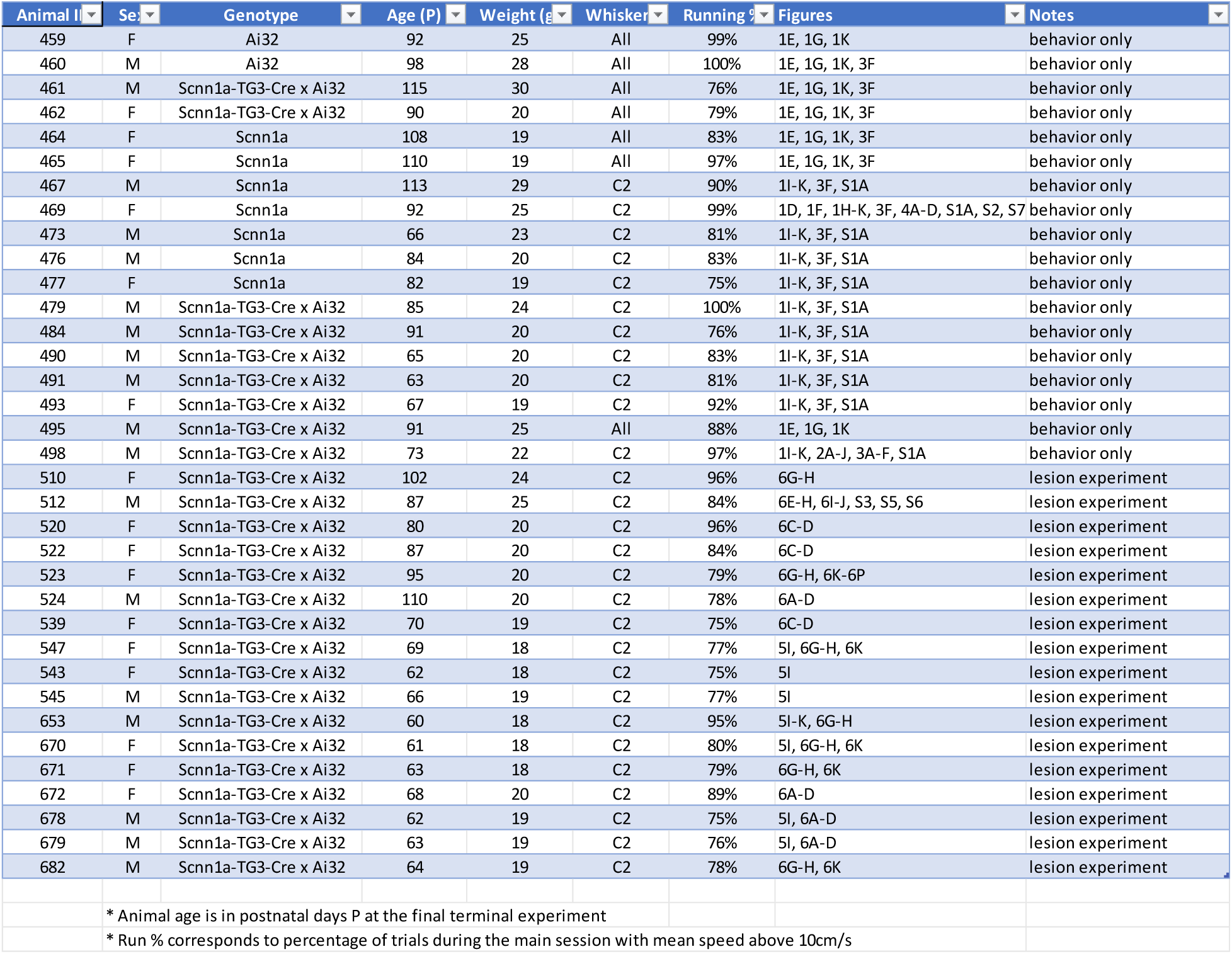
Experimental datasets. Details of animals and which figures they are used in.

| Animal ID | Sex | Genotype | Age (P) | Weight (g) | Whisker | Running | Figures | Notes |
| --- | --- | --- | --- | --- | --- | --- | --- | --- |
| 459 | F | Ai32 | 92 | 25 | All | 99% | 1E, 1G, 1K | behavior only |
| 460 | M | Ai32 | 98 | 28 | All | 100% | 1E, 1G, 1K, 3F | behavior only |
| 461 | M | Scnn1a-TG3-Cre x Ai32 | 115 | 30 | All | 76% | 1E, 1G, 1K, 3F | behavior only |
| 462 | F | Scnn1a-TG3-Cre x Ai32 | 90 | 20 | All | 79% | 1E, 1G, 1K, 3F | behavior only |
| 464 | F | Scnn1a | 108 | 19 | All | 83% | 1E, 1G, 1K, 3F | behavior only |
| 465 | F | Scnn1a | 110 | 19 | All | 97% | 1E, 1G, 1K, 3F | behavior only |
| 467 | M | Scnn1a | 113 | 29 | C2 | 90% | 1I-K, 3F, S1A | behavior only |
| 469 | F | Scnn1a | 92 | 25 | C2 | 99% | 1D, 1F, 1H-K, 3F, 4A-D, S1A, S2, S7 | behavior only |
| 473 | M | Scnn1a | 66 | 23 | C2 | 81% | 1I-K, 3F, S1A | behavior only |
| 476 | M | Scnn1a | 84 | 20 | C2 | 83% | 1I-K, 3F, S1A | behavior only |
| 477 | F | Scnn1a | 82 | 19 | C2 | 75% | 1I-K, 3F, S1A | behavior only |
| 479 | M | Scnn1a-TG3-Cre x Ai32 | 85 | 24 | C2 | 100% | 1I-K, 3F, S1A | behavior only |
| 484 | M | Scnn1a-TG3-Cre x Ai32 | 91 | 20 | C2 | 76% | 1I-K, 3F, S1A | behavior only |
| 490 | M | Scnn1a-TG3-Cre x Ai32 | 65 | 20 | C2 | 83% | 1I-K, 3F, S1A | behavior only |
| 491 | M | Scnn1a-TG3-Cre x Ai32 | 63 | 20 | C2 | 81% | 1I-K, 3F, S1A | behavior only |
| 493 | F | Scnn1a-TG3-Cre x Ai32 | 67 | 19 | C2 | 92% | 1I-K, 3F, S1A | behavior only |
| 495 | M | Scnn1a-TG3-Cre x Ai32 | 91 | 25 | All | 88% | 1E, 1G, 1K | behavior only |
| 498 | M | Scnn1a-TG3-Cre x Ai32 | 73 | 22 | C2 | 97% | 1I-K, 2A-J, 3A-F, S1A | behavior only |
| 510 | F | Scnn1a-TG3-Cre x Ai32 | 102 | 24 | C2 | 96% | 6G-H | lesion experiment |
| 512 | M | Scnn1a-TG3-Cre x Ai32 | 87 | 25 | C2 | 84% | 6E-H, 6I-J, S3, S5, S6 | lesion experiment |
| 520 | F | Scnn1a-TG3-Cre x Ai32 | 80 | 20 | C2 | 96% | 6C-D | lesion experiment |
| 522 | F | Scnn1a-TG3-Cre x Ai32 | 87 | 20 | C2 | 84% | 6C-D | lesion experiment |
| 523 | F | Scnn1a-TG3-Cre x Ai32 | 95 | 20 | C2 | 79% | 6G-H, 6K-6P | lesion experiment |
| 524 | M | Scnn1a-TG3-Cre x Ai32 | 110 | 20 | C2 | 78% | 6A-D | lesion experiment |
| 539 | F | Scnn1a-TG3-Cre x Ai32 | 70 | 19 | C2 | 75% | 6C-D | lesion experiment |
| 547 | F | Scnn1a-TG3-Cre x Ai32 | 69 | 18 | C2 | 77% | 5I, 6G-H, 6K | lesion experiment |
| 543 | F | Scnn1a-TG3-Cre x Ai32 | 62 | 18 | C2 | 75% | 5I | lesion experiment |
| 545 | M | Scnn1a-TG3-Cre x Ai32 | 66 | 19 | C2 | 77% | 5I | lesion experiment |
| 653 | M | Scnn1a-TG3-Cre x Ai32 | 60 | 18 | C2 | 95% | 5I-K, 6G-H | lesion experiment |
| 670 | F | Scnn1a-TG3-Cre x Ai32 | 61 | 18 | C2 | 80% | 5I, 6G-H, 6K | lesion experiment |
| 671 | F | Scnn1a-TG3-Cre x Ai32 | 63 | 18 | C2 | 79% | 6G-H, 6K | lesion experiment |
| 672 | F | Scnn1a-TG3-Cre x Ai32 | 68 | 20 | C2 | 89% | 6A-D | lesion experiment |
| 678 | M | Scnn1a-TG3-Cre x Ai32 | 62 | 19 | C2 | 75% | 5I, 6A-D | lesion experiment |
| 679 | M | Scnn1a-TG3-Cre x Ai32 | 63 | 19 | C2 | 76% | 5I, 6A-D | lesion experiment |
| 682 | M | Scnn1a-TG3-Cre x Ai32 | 64 | 19 | C2 | 78% | 6G-H, 6K | lesion experiment |
| * Animal age is in postnatal days P at the final terminal experiment |  |  |  |  |  |  |  |  |
| * Run % corresponds to percentage of trials during the main session with mean speed above 10cm/s |  |  |  |  |  |  |  |  |

**Figure S1.**
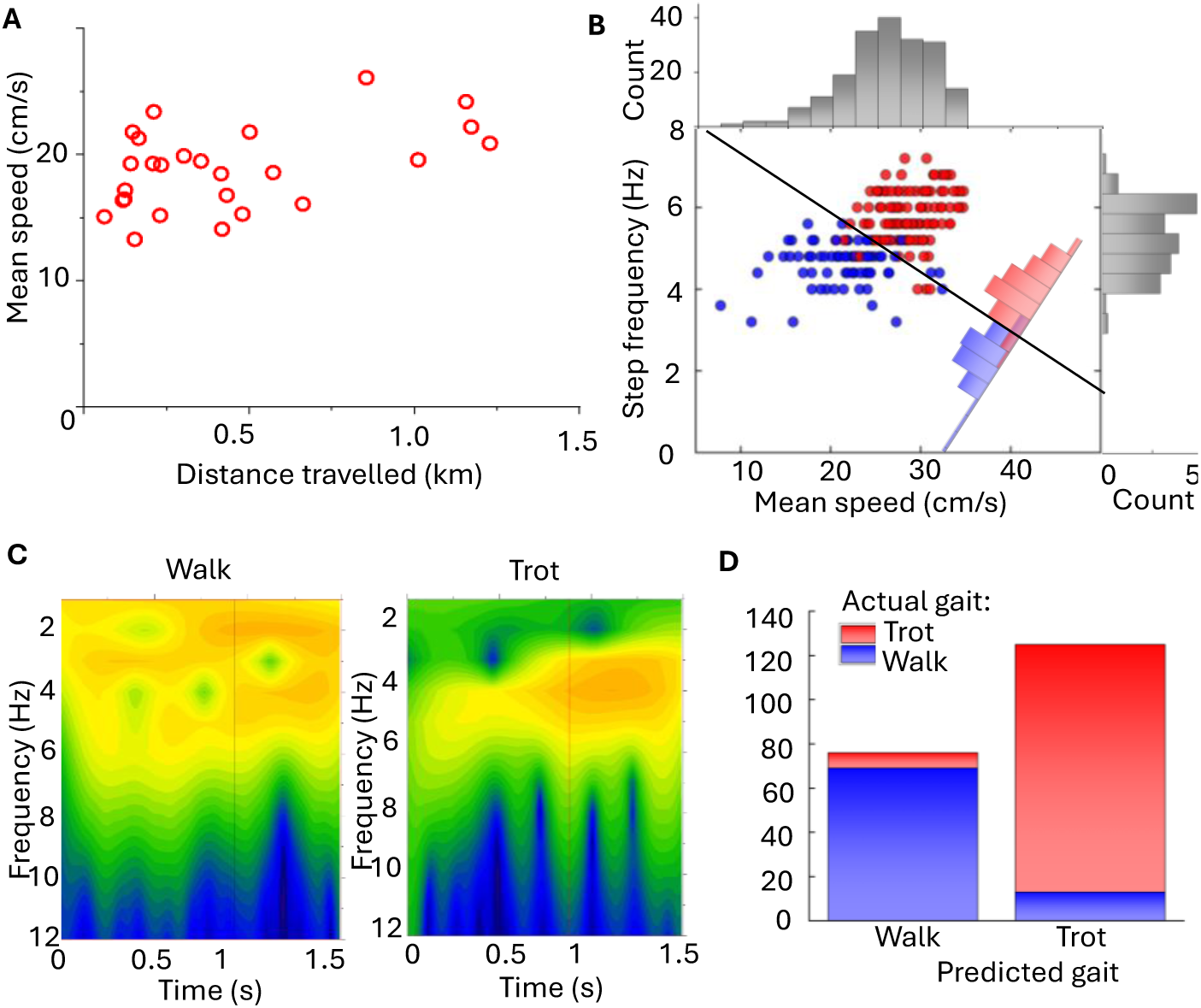
Linear discriminant analysis on locomotion for classification of gaits. **(a)** Mean speeds and total distance travelled for 17 mice during continuous unrewarded navigation session. **(b)** Example Fisher’s LDA results for one animal, 194 total trials, with 118 trotting trials (red) and 76 walking trials (blue) confirmed from manual inspection of lateral acceleration trace and video. Gaits are indistinguishable from mean speeds or step frequency alone. **(c)** Wavelet analysis on lateral acceleration trace from 1.5 second segments of walking and trotting, showing presence of lower frequencies (< 4 Hz) in walking only. **(d)** The results for an example animal with manual inspection to confirm which trials are erroneously assigned. Only trials with a probability of being either walking or trotting >0.8 were used in main figures. Misclassified gait trials are typically due to both gaits being present during the same trial.

**Figure S2.**
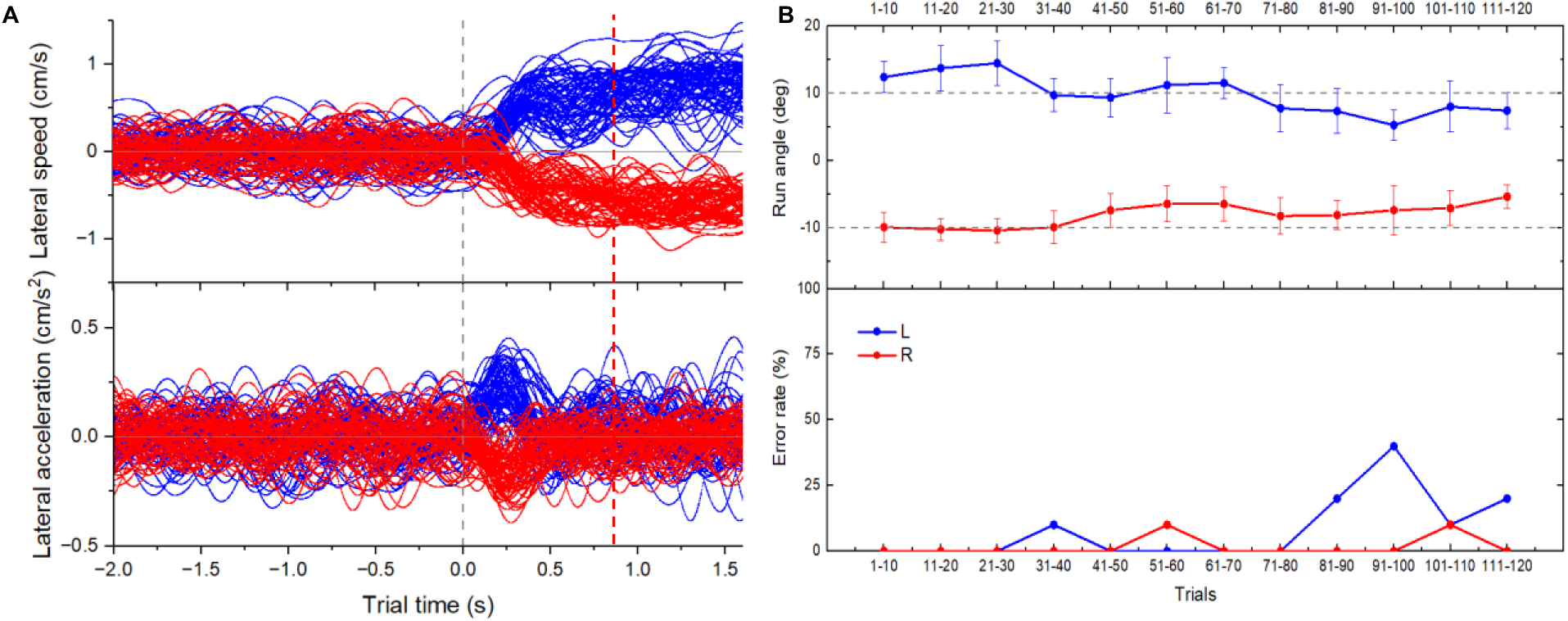
Behavioral performance across main session. **(a)** Main session for an example animal showing trajectories for all OL left trials (blue) and OL right trials (red) overlaid. Lateral speed (top) and lateral acceleration (bottom) traces show animal clearly distinguishing between turn types with consistent peak in lateral acceleration that corresponds to RT. **(b)** Performance over the session for the same animal. Run angle (top) shows mean and standard deviation of run angles during the turning trial for groups of 10 trials in chronological order over a single main session. Even though the angle itself may be below 10°, errors (bottom) are defined as a change in mean run angle < 10° between turning trial and preceding CL straight trial.

**Figure S3.**
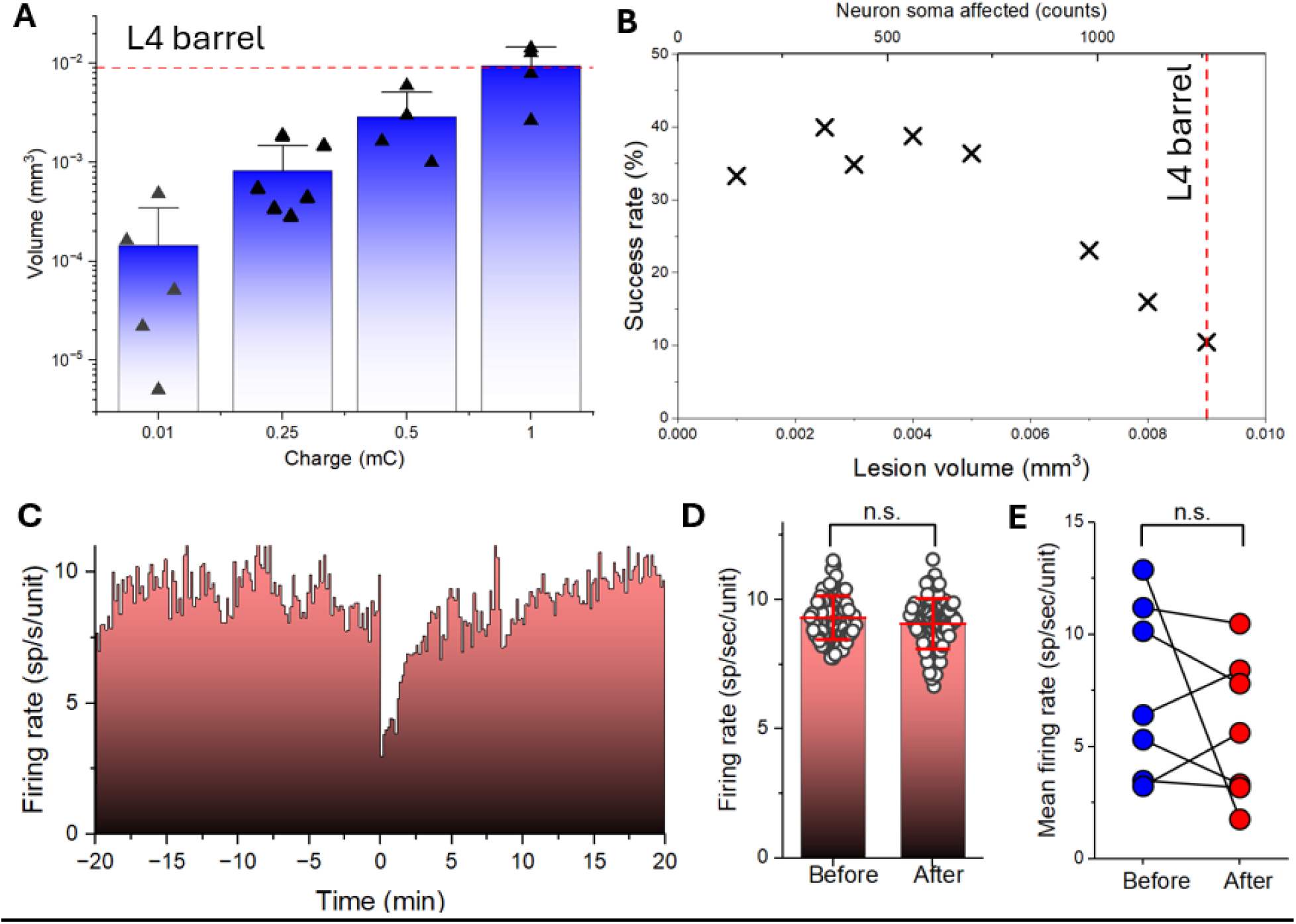
Lesion calibration. **A)** A series of experiments were carried out to calibrate the charge provided to the lesion electrode and subsequent volume of the resulting lesion (n = 3). Current of up to 500µA was applied for application time up to 2 seconds to produce charge in the range of 0.01mC to 1mC. **B)** Degradation of success rate in contralateral trials for different lesion volumes produced in principal C2 barrel. **C)** Normalized firing rate for C2 barrel column units before (negative times) and after (positive times) lesion of 10% of L4 barrel volume is produced (45 units pre-lesion, 33 units post-lesion). **D)** Statistics of firing rates before (left bar) and after (right bar) lesion of 10% of L4 barrel volume is produced for the same animal as in C). **E)** Mean normalized firing rates before (blue circles) and after (red circles) lesions are produced in the principal C2 barrel for n=7 animals.

**Figure S4.**
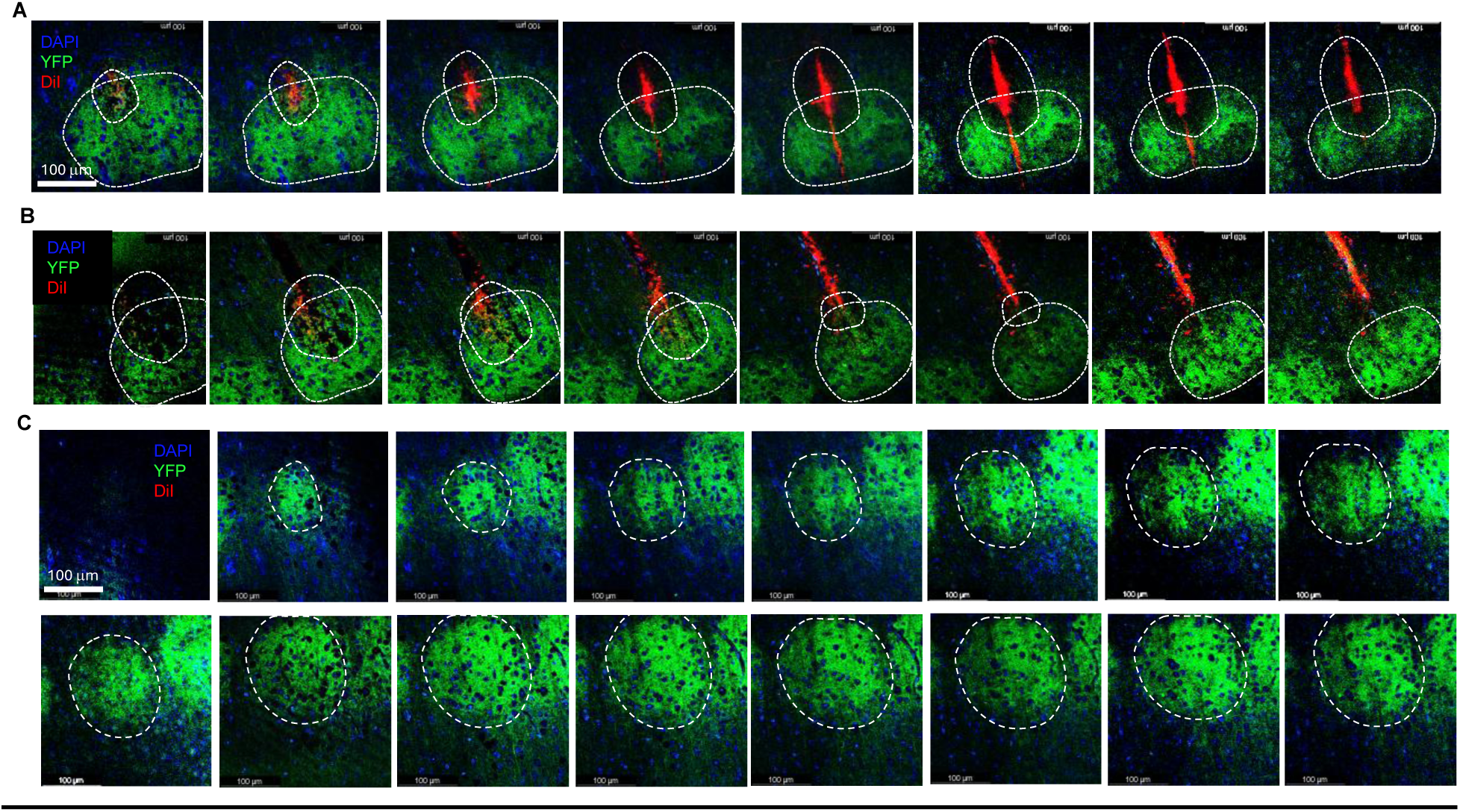
Z-stack image series of a lesion in C2 barrel for cell counting and quantification. **(a)** Lesion in contra-lateral column. 8 images of an example lesion coronal slice covering total 40µm depth of slice section with 5µm separation between each image. 50 Z-stack images total were taken using confocal microscope for each 50µm slice with 1µm separation in full series, used for volumetric reconstruction. Each image is an overlay of EYFP (green), DiI (red) and DAPI (blue) fluorescence images. **(b)** 8 images of the next 50µm slice of the same contra-lateral lesion in the same animal covering total 40µm depth with 5µm separation between each image. **(c)** Representative 24 images for full Z stack of ipsilateral un-lesioned C2 barrel for cell count and quantification comparison with the lesioned barrel in Fig. S5. Each image is 5µm apart in depth. Each image is an overlay of EYFP (green), DiI (red) and DAPI (blue) fluorescence images.

**Figure S5.**
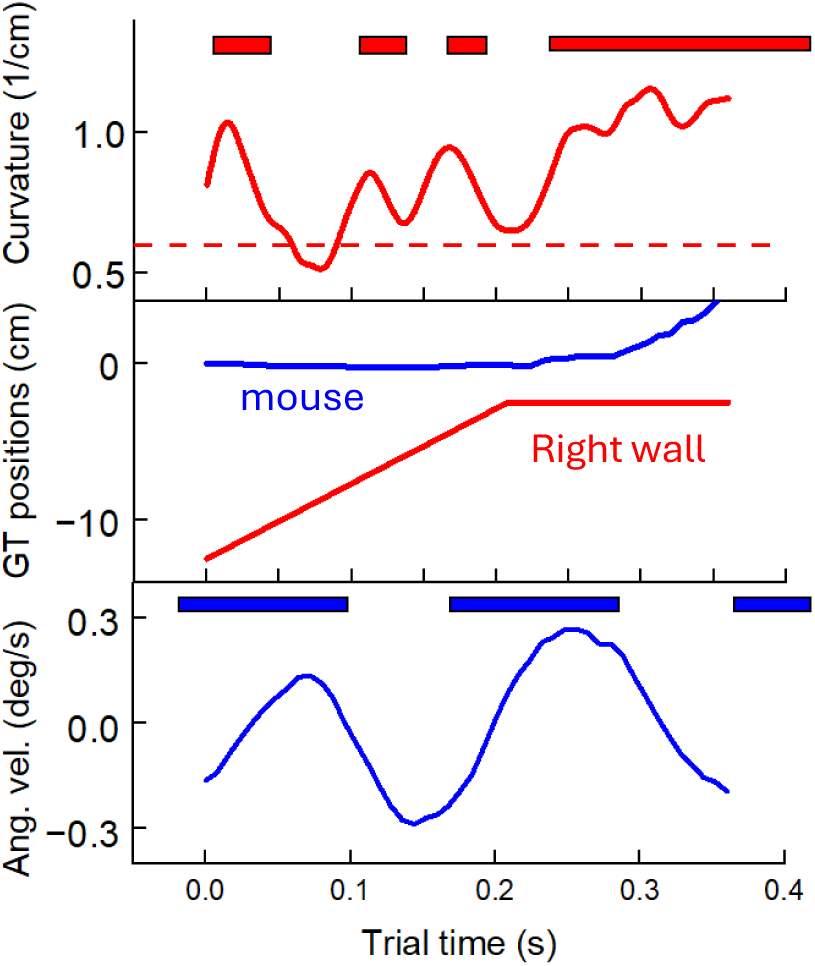
Whisker and locomotion variables, for example CL trial. Example section of a CL trial used in Fig.6E. Top: whisker contact times with the wall (red rectangles) and whisker curvature extracted from overhead video. Middle: Positions of the right wall (red) and trajectory of the mouse (blue) used as ground truth (GT) for Fig.6E. Bottom: RH leg contact times with the sphere (blue rectangles). Angular velocity trace showing characteristic maximum at the maximum propulsion by RH leg to produce a turn.

**Movie S1.**
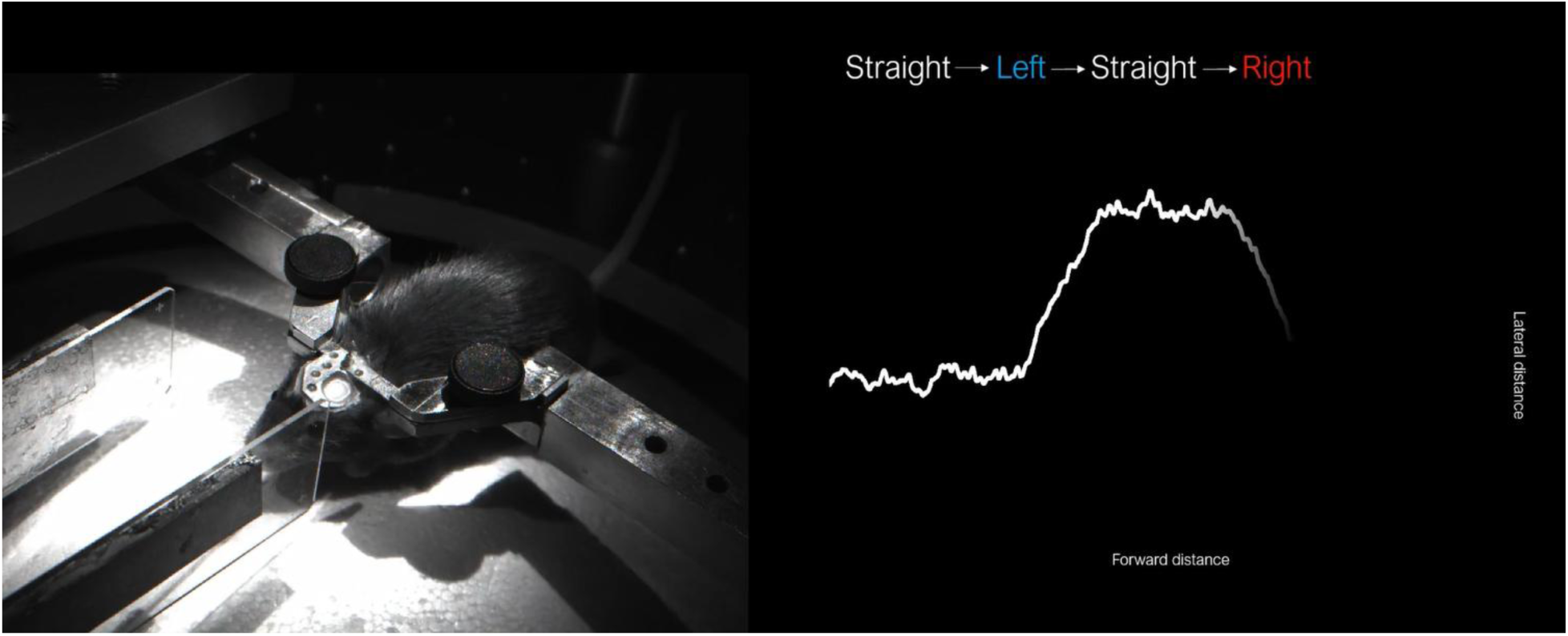
Left: Movie of two consecutive trials. Right: Trajectory of a mouse during these trials. Each trial consists of two parts (Fig.4A, top): a section of straight CL wall tracking for 100cm forward distance as in the habituation sessions, followed by a 2 second-long open loop (OL) section when the CL feedback is terminated, and the walls are first briefly removed beyond the reach of the C2 whiskers and then just one of the walls rapidly approaches the snout (within 0.2s) and stays at a pre-defined constant distance for 1.8 seconds before it retracts again. First trial is a left turn when the right wall is brought in closer to the animal. The mouse senses the approaching wall with its right C2 whisker and is forced to turn either to the left to avoid the expected collision. Second trial is a right turn when left wall is approaching. Randomized sequence of these left and right turn trials creates a version of an ANC task

